# Cancer-Associated Mesothelial Cells Drive Immune Escape and Therapy Resistance in Ovarian Cancer

**DOI:** 10.64898/2026.01.07.698232

**Authors:** Maeva Chauvin, Julien Roche-Prellezo, Virginie Lafont, Henri-Alexandre Michaud, Estelle Tromelin, Robin Michel, Clara Freixinos, Marie-Charlotte Meinsohn, Pierre-Emmanuel Colombo, Nathalie Bonnefoy, Laurent Gros, David Pépin

**Affiliations:** Pediatric Surgical Research Laboratories, Massachusetts General Hospital, Boston, MA, USA - Department of Surgery, Harvard Medical School, Boston, MA, USA; IRCM, Univ Montpellier, ICM, INSERM, Montpellier, France; Plateforme de Cytométrie et d’Imagerie de Masse de Montpellier, Institut de Recherche en Cancérologie de Montpellier, INSERM U1194 - UM – ICM, Montpellier, France; Department of Obstetrics and Gynecology, University of Rennes, Rennes, France

## Abstract

Cancer-associated mesothelial cells (CAMCs) are key modulators of the ovarian tumor microenvironment, contributing to tumor growth an immune evasion. Normal mesothelial cells play a role in peritoneal homeostasis and immune surveillance and represent the first point of contact during abdominal dissemination of ovarian cancers. Yet, their role in ovarian tumor immunity remains poorly understood. Here, we map the cellular states, spatial organization, and immune functions of CAMCs across ovarian cancer progression. Using lineage tracing and spatial transcriptomics, we demonstrate that CAMCs originate from mesothelial cells at the tumor surface and can progressively infiltrate the tumor core, undergoing a phenotypic transition towards fibroblast-like and immunosuppressive states. We characterize the function of an unrecognized CAMC subtype marked by SERPINB2^+^ expression, and a combination of markers absent in normal mesothelial cells. CAMC^Serpinb2+^ cells have reduced expression of pro-inflammatory cytokines (IL-2, IL-7, IL-12, IL-15) and increased expression of IL-10, TGFß1, and CCL17, promoting regulatory T cell recruitment and tolerogenic CD4^+^ T cell responses. Functionally, CAMC^Serpinb2+^ accelerate tumor growth, reduce CD4^+^ T and B cell infiltration, and expand Treg populations, ultimately leading to immunotherapy resistance. Together, our findings identify CAMCs as a potential therapeutic target in peritoneal carcinomatosis and as a means of restoring sensitivity to current treatments.

**Graphical abstract:** 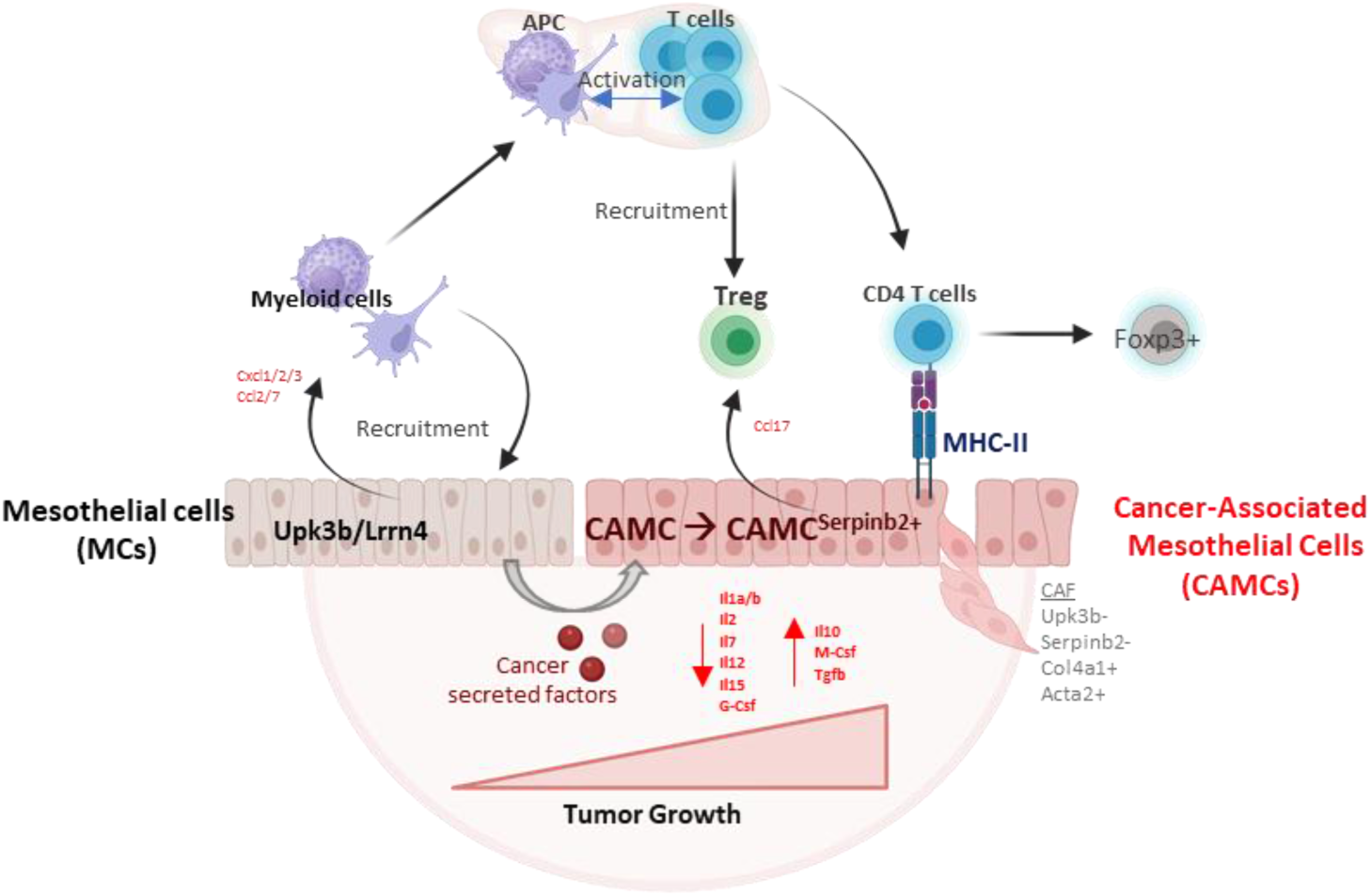

## INTRODUCTION

High-grade serous ovarian cancer (HGSOC) remains one of the most lethal of the gynecological malignancies, which is characterized by late-stage diagnosis, rapid progression, and results in 70% recurrence within two years^1–3^. Despite advancements in targeted therapies and immunotherapies, the intricate interplay between cancer cells and the tumor microenvironment (TME) continues to pose significant challenges in effectively managing HGSOC. Among the various components of the TME, cancer-associated mesothelial cells (CAMCs) have emerged as pivotal players, influencing tumor behavior and immune responses^4,5^.

Mesothelial cells (MCs), which line the peritoneal cavity, have long been recognized for their role in maintaining serosal integrity and facilitating tissue homeostasis^6–8^. However, during tumor progression, these cells undergo significant alterations, acquiring characteristics associated with tumor promotion and immune evasion^4^. During metastasis to the omentum, cancer cells are thought to attach to the mesothelium and invade the omental tissue, and in that process reprogram mesothelial cells into cancer-associated mesothelial cells (CAMCs) that integrate into the tumor stroma and promote cancer growth^5,9–13^. CAMCs have been described to regulate many cell types of the TME including macrophages and cancer-associated-fibroblasts (CAFs), by secreting growth factors, cytokines, and chemokines ^10,14,15^. Their immunomodulatory function is reminiscent of the role of normal peritoneal mesothelium in protecting peritoneal organs by producing serosal fluids and surveying the cavity for pathogens to activate innate and adaptive immune responses at the serosal surface in case of infection^16,17^. Mesothelial cells express multiple cytokines, antimicrobial peptides, and growth factors that, in turn, regulate the proliferation and differentiation of immune cells, including T cells^18^. Similarly, in tumors, the secretion of TGFb by ovarian cancer cells induces the production of cytokines by CAMCs, which, in turn, regulate cytotoxic T lymphocytes^19^. This paracrine effect can modulate response to anti-PD-1 therapy and contribute to resistance to immunotherapies, particularly in ovarian cancer^6,20,21^. Yet the role of CAMCs in regulating growth and immunosuppression in ovarian cancer remains poorly understood.

In this study, we characterized CAMCs subtypes and cell states present in TME of ovarian cancer and begin to explore their underlying function, including a potential immunosuppressive phenotype that may contribute to immune evasion. We speculate that therapeutic strategies eliminating or reprogramming CAMCs could restore a more immunogenic TME and a better response to immunotherapy.

## RESULTS

### Tumor stroma contains different subtypes of CAMCs in a syngeneic mouse model of ovarian cancer

To define the gene signature of cancer-associated mesothelial cells (CAMCs) in the tumor microenvironment (TME), we analyzed single-cell RNA sequencing (scRNA-seq) datasets from omental tumors of a syngeneic ovarian cancer mouse model (KPCA, GSE233423)^22^. Briefly, this transcriptome was derived from applying the 10X scRNAseq pipeline to dissociated omental metastases generated by grafting the KPCA (displaying **K**RAS, **P**TEN, **C**CNE1, **A**KT2 mutations) transformed fallopian tube epithelium cell line into the peritoneal cavity of syngeneic C57BL/6 mice, to mimic human CCNE1-amplified HGSOC, as previously described^13,22^ (Fig.1A). CAMCs were identified by expression of the classical mesothelial markers *Upk3b* and *Lrrn4*^13,23,24^, and were distinguishable from CAFs, which expressed classical fibroblast markers such as *Acta2* and *Tgln*^25^ (Fig. S1A). Subclustering analysis of the CAMCs in the mouse tumors revealed three subsets of CAMCs: 1) a proliferative CAMC cluster marked by mitotic genes such as *Top2a*, *Ccne2*, *Prc1*, and *mKi67*^26^, and two clusters of CAMC with distinct transcriptional signatures, defined by Serpinb2 marker expression as 2) CAMC^serpinb2-^, and 3) CAMC^serpinb2+^ (Fig. 1A/B). Indeed, when we examined the top 10 highly expressed genes of CAMC ^Serpinb2+^ cells (*Serpinb2*, *Ero1l*, *Asn*s, *Adm2*, *Aldh1l2*, *Mthfd2*, *Phgdh*, *Vegfa*, *Amhr2*.) across all cell populations in the tumor microenvironment, we found that Serpinb2 was mostly absent in other cell types, and specific to this subpopulation of CAMCs (Fig. 1E). The CAMC^serpinb2+^ gene expression profile revealed by scRNAseq was significantly enriched for gene ontology terms related to metabolism, such as the L-serine biosynthetic process (Fig. S1C). The CAMC^Serpinb2-^ cluster was significantly enriched for gene ontology terms related to positive regulation of type 2 immune response and antigen presenting via MHCII and associated with high level of MHC-II subunits (*H2-Aa*, *H2-Eb1*, *H2-Ab1*, *Cd74*) (Fig. 1B, Fig. S1A). We hypothesize that all three CAMC populations arose from normal peritoneal mesothelial cells (*Upk3b*+/*Lrrn4*+) responding to the cancer microenvironment. These populations and their differential gene expression are highlighted in a heatmap (Fig. S1A) and dot plot (Fig. 1B).

**Figure 1:**
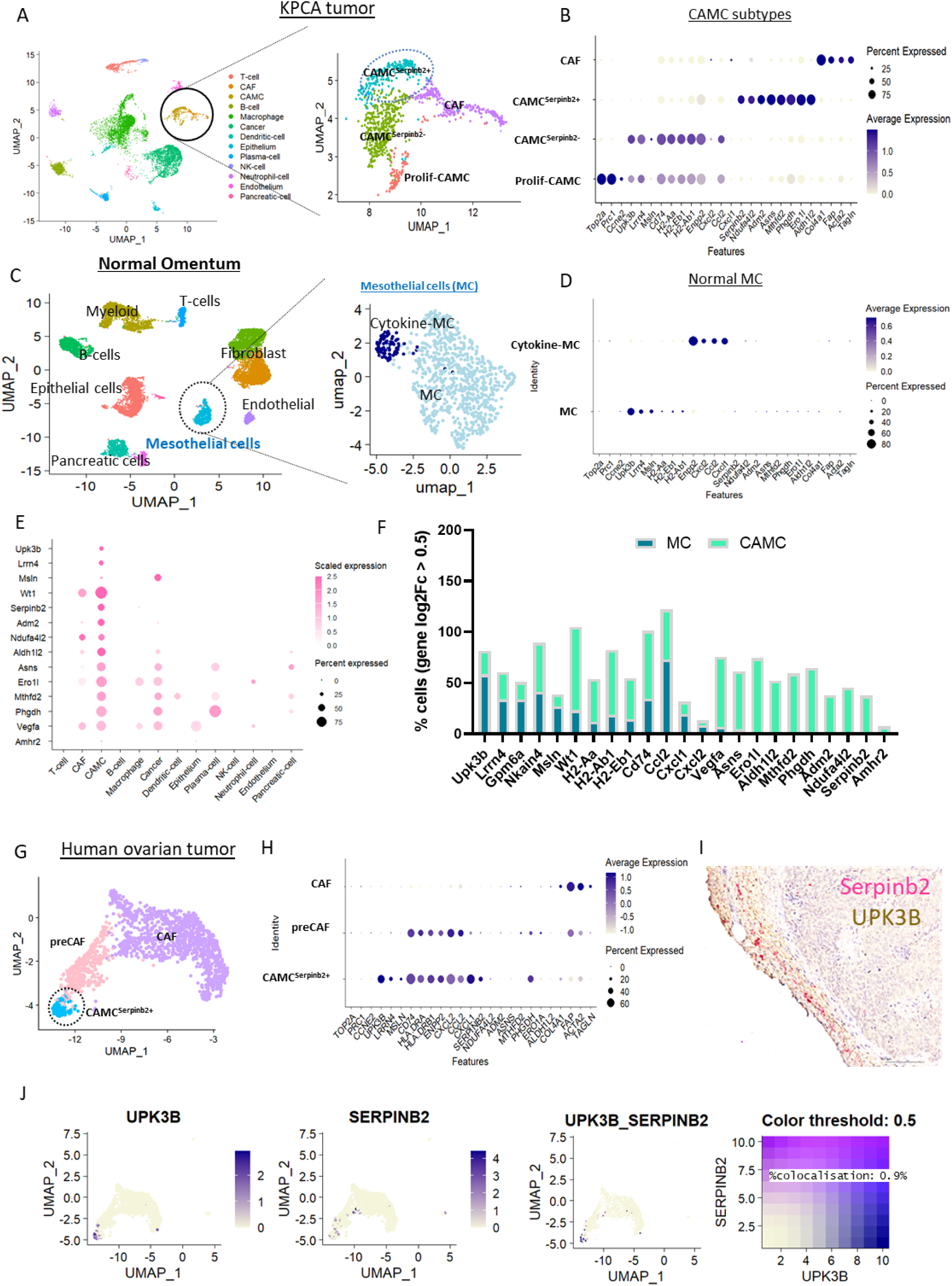
Identification of CAMC subtype gene signatures in ovarian cancer microenvironment. (A) Right panel: UMAP representation of single-cell RNA-seq (scRNA-seq) data from KPCA tumors (GSE233423), showing the cellular composition of the tumor microenvironment. Left panel: UMAP focused on the stromal compartment, highlighting cancer-associated mesothelial cell (CAMC) subtypes (Prolif-CAMC, CAMCSerpinb2-, CAMCSerpinb2+) and cancer-associated fibroblast (CAF) clusters. (B) Dot plot showing the percentage and average expression levels of CAMC subtype–specific genes among the top 50 genes. Dot size indicates the proportion of cells expressing each gene, while color intensity represents average expression (dark blue = high expression, beige = low expression). (C) UMAP of cell populations from the omentum and UMAP of mesothelial cell (MC) clusters extracted from scRNA-seq data (GSE134355). (D) Dot plot of CAMC subtype–specific gene expression across MC clusters. (E) Dot plot of CAMC marker expression across all KPCA tumor cells. (F) Relative proportion of cells expressing CAMC subtype gene signatures compared to normal mouse primary MCs (gene expression: Log_2_FC > 0.5). (G) UMAP highlighting human CAMC subtypes and CAF clusters from ovarian tumors (GSE147082). (H) Dot plot showing gene expression levels of CAMC and CAF subtypes in humpan ovarian tumor. (I) Representative high-grade serous ovarian cancer (HGSOC) tissue section showing *SERPINB2* expression detected by in situ hybridization (pink dots) and UPK3B expression by immunohistochemistry (brown staining), demonstrating spatial co-localization at the tumor–stromal interface. (J) Feature plot showing *SERPINB2* and *UPK3B* expression in human CAMC and CAF subtypes and

### The CAMC^Serpinb2+^ transcriptional signature is unique to the tumoral context

To characterize the pathogenic states of CAMCs signatures within the tumor microenvironment, we aimed to contrast their transcriptomic profiles to normal mesothelial cells (MCs) extracted from mouse omenta. We took advantage of scRNA-seq datasets (GSE134355)^27^ that included cell populations from healthy mice omenta (N=4) (Fig. 1C). Clustering of MCs (*Upk3b*+/*Lrrn4*+) revealed a distinct subset enriched in CXCL/CCL chemokines (Cytokine-MCs: *Ccl2*, *Cxcl1/2*) (Fig. 1C/D), potentially indicative of an activated immune state^28^. We then examined the proportion of cells in normal omental MCs expressing genes specifically associated with the CAMC ^Serpinb2-^ and CAMC ^Serpinb2+^ (Log2FC > 0.5) pathogenic states found in the TME (Fig. 1F). *Upk3b*, *Lrrn4*, *Gpm6a*, *Nkain4*, *Msln*, and MHC-II subunits (*H2-Ab*, *H2-Ab1*, *H2-Eb1*, *Cd74*) were detected in both normal MCs and pathogenic CAMC^Serpinb2-^ and CAMC^Serpinb2+^ populations. However, MHC-II subunit expression was significantly enriched in CAMCs, suggesting a heightened antigen-presenting function (Fig.1F). Interestingly, markers of CAMC^Serpinb2+^ cells (*Serpinb2*, *Adm2*, *Ndufa4l2*, *Asns*, *Aldh1l2*, *Mthfd2*, *Phgdh*, *Vegfa*, *Amhr2*) were unique to the tumor context. Based on this gene expression data, we decided to further investigate the immune functions of CAMCs, and particularly the role CAMC^Serpinb2+^ on tumor immunity.

We next investigated whether CAMC subtype signatures were conserved in human HGSOC tumors. Analysis of the untreated HGSOC patient dataset (GSE147082, n=6) ^29^ revealed a CAMC^SERPINB2+^ cluster in the human TME. In contrast to KPCA tumors, HGSOC samples lacked well-defined *SERPINB2*+ and *SERPINB2*-populations, perhaps due to the limited amount of CAMCs represented in this dataset. *SERPINB2* and *UPK3B* were largely expressed within the same cluster (Fig. 1G/I), although SERPINB2 showed broader distribution, extending into the pre-CAF cluster, and the proportion of double-positive cells (UPK3B+/SERPINB2+) was very low (0.9%), with limited colocalization (Fig. 1J, 1I), suggesting these phenotypes are not overlapping. We hypothesize that the CAMC^Serpinb2+^ cells arise from the differentiation of UPK3B+ SERPINB2- cells. Similarly, to the mouse, the CAF cluster appeared transcriptionally distinct from CAMCs, and distinguishable by the expression of fibroblast markers such as ACTA2, COL4A1, FAP, and TAGLN.

### Normal mesothelial cells acquire a CAMC^Serpinb2+^ signature in a time-dependent manner following stimulation by cancer-secreted factors

To understand the dynamic changes occurring during the differentiation of MCs into CAMC subtypes, as defined by expression of cluster-specific markers such as *Upk3b*, *H2-Aa* (MHCII subunit), *Serpinb2*, and *Col4a1* (Fig. 2A), we predicted their developmental trajectories based on gene expression dynamics over pseudotime (Fig. 2B). Pseudotime analysis suggested a linear transition from the CAMC ^Serpinb2-^ to the CAMC^Serpinb2+^ state, observed both in murine CAMCs (top panel) and in human HGSOC (bottom panel) (Fig. 2B).

**Figure 2:**
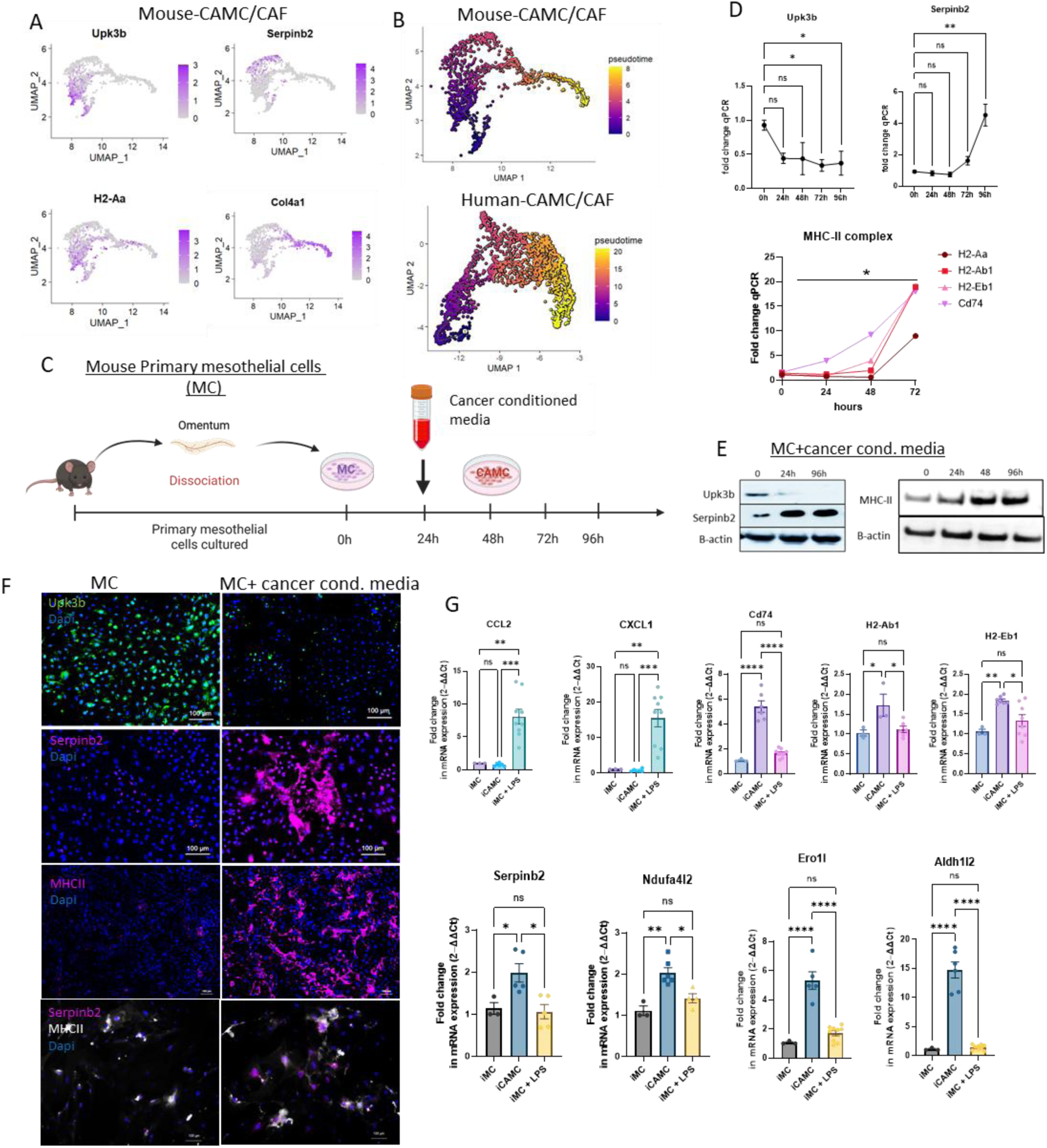
Differentiation of Mesothelial cells into CAMC^Serpinb2+^ overtime. A) FeaturePlot of *Upk3b, Serpinb2, H2-Aa* and *Col4a1* expression to represent CAMC and CAF subtypes. B) Pseudotime analysis of single-cell RNA sequencing (scRNA-seq) of KPCA tumors and human ovarian cancer. The color gradient reflects the progression of pseudotime, with lighter colors (yellow) indicating early stages and darker colors (darkblue) representing later stages. Genes of interest are plotted along the pseudotime axis, illustrating their changes in expression levels throughout the differentiation process. C) Illustration of the experimental design showing primary mesothelial cells harvested from murine omentum and reprogrammed into CAMCs by stimulation with cancer-conditioned media over time (0 h, 24 h, 48 h, 72 h, 96 h). D) qPCR analysis of *Upk3b, Serpinb2* and MHCII chains gene expression levels of primary mesothelial cell (MC) stimulated by cancer conditioned media (KPCA) at different time points (0h, 24h, 48h, 72h and 96h). The relative expression of *Upk3b* and *Serpinb2* is shown, normalized to a housekeeping gene (*Gapdh*) and presented as fold change compared to the 0-hour time point. Data are presented as mean ± SEM (n = 3); statistical significance is indicated by *p < 0.05 and **p < 0.01 compared to the 0-hour time point (t-test). E) Western blot analysis of UPK3B, SERPINB2 and MHCII protein levels in CAMC ranging to 0h-96hrs. Representative blots show the relative abundance of Upk3b and Serpinb2 proteins, with β-actin used as a loading control. F) Immunofluorescence staining of UPK3B and SERPINB2 of mesothelial cells (MC) stimulated by cancer conditioned media at 72hrs. Representative images show UPK3B, SERPINB2 and MhcII protein expression, with DAPI used for nuclear staining. Scale bar: 100 μm. Fluorescence intensity quantification is presented as mean ± SEM (n = 3); significance is indicated by *p < 0.05, **p < 0.01 (t-test). G) Expression levels of representative CAMC^Serpinb2+^ genes (*Serpinb2, Ndufa4l2, Ero1l, Aldh1l2*), MHC subunits (*Cd74, H2-Ab1*) and cytokines (*Cxcl1, Ccl2*), measured by qPCR in immortalized mesothelial cells (iMC) stimulated with either KPCA-conditioned media or 1 ug/ml lipopolysaccharide (LPS).

To better understand how normal mesothelial cells may recapitulate this differentiation into the CAMC^Serpinb2+^ state, we examined transcriptional changes in primary MCs, isolated from normal omenta, as they differentiated into CAMCs following stimulation with media conditioned by KPCA cancer cells (24h conditioning). We then followed the expression of CAMC^Serpinb2+^ markers in a time-dependent manner (0 – 96 hrs) (Fig. 2C), assessing *Upk3b*, MHCII sub-units, and *Serpinb2* expression by qPCR (Fig. 2D). We observed a significant decrease in *Upk3b* expression after 24 hours of cancer media stimulation, while MHCII subunits and *Serpinb2* expression was increased significantly at 72 and 96 hours, respectively. To decipher the full transcriptomic changes associated with MC differentiation into CAMCs we performed bulk RNA-seq following 72 hours of stimulation with KPCA cancer-conditioned media. This analysis consistently identified an increase in Serpinb2, Ndufa4l2, Ero1a, and other markers of CAMC^Serpinb2+^ cells found in KPCA tumors, as well as an increase in MHC-II subunits (*H2-Ab1*) generally observed in CAMCs (Fig. S2A/B). Notably, geneset enrichment analysis (GSEA) of differentially expressed genes (Fig. S2B) revealed an enrichment of pathways related to T-cells, cytokines, and metabolism (Fig. S2C).

We further validated the transcriptional changes in *Upk3b*, MHCII subunits, and *Serpinb2* at the protein level in primary MCs stimulated by KPCA media after 24 and 96 hours of stimulation by Western blot (Fig. 2E). Consistent with the qPCR data, we observed a reduction in UPK3B and an induction of MHCII and SERPINB2 when compared to unstimulated normal mesothelial cells (Fig. 2E). Similar findings were confirmed by immunofluorescence analysis at 72 hours (Fig. 2F). Immunofluorescence analysis of MHCII and SERPINB2 staining showed extensive co-localization in primary MCs following stimulation with KPCA-conditioned media, suggesting that the CAMC^Serpinb2+^ arise through a linear transition from MCs MHCII+ CAMC^Serpinb2-^ cells, to MHCII+ CAMC^Serpinb2+^ (Fig. 2F).

To reduce batch-to-batch variability of primary MCs, we established an immortalized mesothelial cell line, iMC (SV40T-transformed mouse primary mesothelial cells). We confirmed that iMCs respond similarly to primary MCs when stimulated with cancer cell conditioned media by recapitulating the downregulation of Upk3b expression and upregulation of Serpinb2 expression by qPCR, Western blot, and immunofluorescence (Fig. S3). This pattern was also observed in a human mesothelial cell line (Met5a) following stimulation by primary cancer cells derived from a high-grade serous ovarian cancer patient (HGSOC pt-D)^30^. However, stimulation of Met5a with media conditioned by Ov90 cells, another ovarian cancer cell line, had a more blunted response, only modestly inducing SERPINB2 at the 96-hour timepoint (Fig. S3A/B/C). These results suggest that cancer cells are heterogeneous in their ability to reprogram mesothelial cells into the CAMC^Serpinb2+^ state, which might lead to heterogeneity of CAMC populations across patients.

**Figure 3:**
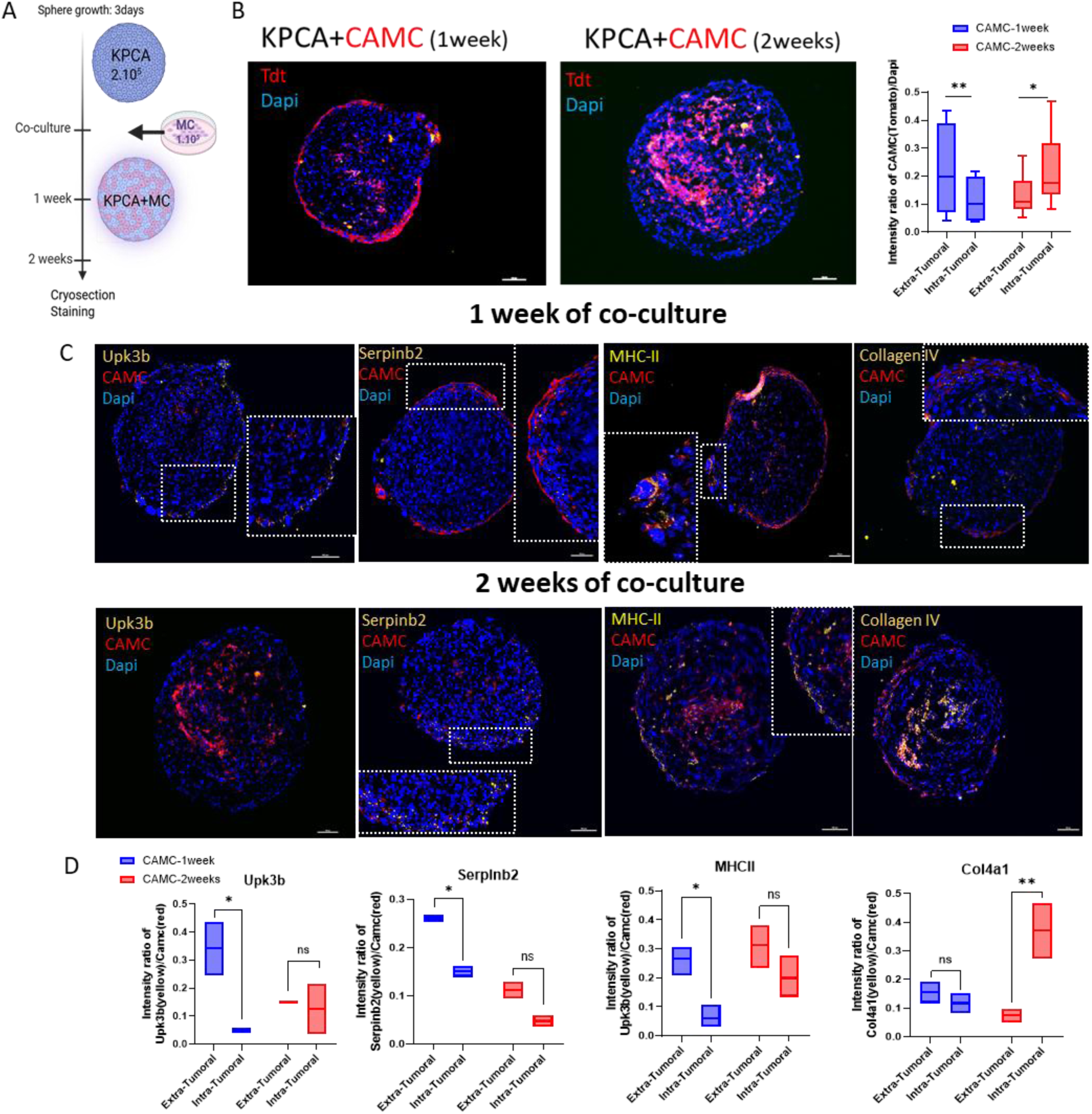
CAMCs Infiltrate the Core of the Spheroid and adopt a CAF-like gene expression signature. A) Illustration of the experimental design for 3D KPCA spheroid formation (2.10^5^ cells). Three days after sphere formation, primary mesothelial cells (tdTomato-reporter) were added at 1.10^5^ cells. Spheres were harvested after 1 and 2 weeks of co-culture with primary mesothelial cells. B) Representative images of 3D spheroids cryosections following 1 and 2 weeks of culture. Percentage of infiltration of MCs quantified at the surface (surface delimited by 50µm contouring) and into tumor. Scale bars 100µm C) Representative micrographs of Immunofluorescence staining of the spheres showing the expression of UPK3B, SERPINB2, and COL4A1 (in yellow) at 1 and 2 weeks. Scale bars 100µm. D) Quantification of intensity staining of UPK3B, SERPINB2, MHCII, and COL4A1 at the surface (extra-tumoral) and in the spheroid core (intra-tumoral) at 1 and 2 weeks.

### The CAMC^Serpinb2+^ signature is specifically induced by cancer-secreted factors and not other inflammatory stimuli such as bacterial LPS

We next sought to determine whether the CAMC^Serpinb2+^ gene signature was specifically induced by tumor-derived factors or can also be triggered by other external inflammatory stimuli. We used lipopolysaccharide (LPS), a component of bacterial cell walls, to induce MCs activation *in vitro*. As expected, MCs responded to LPS stimulation (72h incubation) by upregulating Ccl2 and Cxcl1, two chemokines typically induced during inflammatory responses^31^ (Fig. 2G). Notably, these cytokines were also identified in the cytokine-MC cluster, supporting the idea that this subset reflects endogenous immune activation of MCs (Fig. 1C/D). However, LPS did not modulate the expression of CAMC^Serpinb2+^ markers (*Serpinb2*, *Ndufa4l4, Ero1l*, and *Aldh1l2*) or MHCII sub-units (*Cd74*, *H2-Ab1*, *H2-Eb1*) (Fig. 2G). In contrast, conditioned media from KPCA tumor cells (72h incubation) significantly increased the expression of CAMC^serpinb2+^ genes, suggesting that KPCA-derived factors reprogram MCs through distinct, cancer-specific mechanisms (Fig. 2G).

### Loss of CAMC gene expression signature coincides with mesothelial cell mesenchymal transformation into CAFs and invasion into the tumor core

To investigate the spatiotemporal evolution of CAMC subtypes within the TME, we developed a 3D spheroid model composed of KPCA cancer cells (2.10^5^ cells) for3 days before being co-cultured with freshly isolated MCs from tdTomato reporter mice (1.10^5^ cells) (Fig. 3A). The tdTomato enabled real-time tracking of MC-derived cells during spheroid development. After one week in culture, the spheres were cryosectionned for staining and tdTomato+ CAMCs were observed forming a layer at the spheroid surface, while after two weeks they were actively infiltrating the tumor core (Fig. 3B). Notably, CAMCs located at the spheroid periphery surface expressed UPK3B, MHCII, and SERPINB2, as determined by immunofluorescence staining on spheroid sections (Fig. 3C). The surface expression was quantified by contouring the spheroid periphery (approximately 50 μm thickness) and measuring the intensity of the stained area (Fig. 3D). In contrast, infiltrating CAMCs lacked these markers and instead exhibited COL4A1 expression, suggesting a transition toward a stromal fibroblast–like phenotype within the tumor (Fig. 3C/D), as seen in the UMAP of the stromal cluster from scRNAseq of KPCA tumors (Fig. 2A).

To further investigate whether CAMCs may represent a progenitor for a subset of cancer-associated fibroblasts (CAFs), we assessed the expression of *Acta2* and *Col4a1*, two markers enriched in CAF subpopulations identified in both our murine and human scRNA-seq datasets (Fig. S1). Upon stimulation with cancer cell–conditioned media, primary and immortalized MCs exhibited increased protein levels of ACTA2 and COL4A1 after 96 hours, as determined by immunofluorescence staining (Fig. S3C).

### Spatial analysis of metastases finds an association between CAMCs and CD4^+^ T at the tumor periphery

Anti-Müllerian hormone receptor type 2 (AMHR2) has been previously described as a surface marker of CAMCs^13^. We took advantage of this marker to trace the fate of Amhr2+ CAMC cells as they progress from the CAMC^Serpinb2-^ state to CAMC^serpinb2+,^ and ultimately differentiate into CAFs (Fig. 4A). We utilized the AMHR2-CreERT2/mTmG inducible reporter mouse model^13^, which allowed lineage tracing via tamoxifen pulse-chase induction of GFP (and loss of Tomato) in Amhr2+ CAMCs and their descendants *in vivo*. KPCA tumor cells were engrafted intra-peritoneally into this reporter mouse, and tamoxifen (20 mg/kg) was administered 48 hours later to induce conversion of the floxed mTmG allele in AMHR2-CreERT2+ cells (Fig. 4B). GFP+ cells in tumors were assessed after 10 days of growth by cryosection and immunofluorescence analysis (Fig. 4D). We found GFP+ cells at the tumor surface that co-expressed UPK3B, MHCII, and SERPINB2, with a co-expression average of 77%, 83%, and 78% respectively (N=3). Furthermore, GFP+ cells infiltrating deeper into the tumor mass were also observed, which lacked expression of UPK3B and SERPINB2, but showed robust expression of COL4A1 a marker of CAFs (intra-tumoral GFP-co-expression mean: 81%). These results suggest a phenotypic shift of the CAMC lineage into CAF-like cells (Fig. 4C/D), a finding that parallels our results obtained with our 3D spheroid model (Fig. 3C/D).

**Figure 4:**
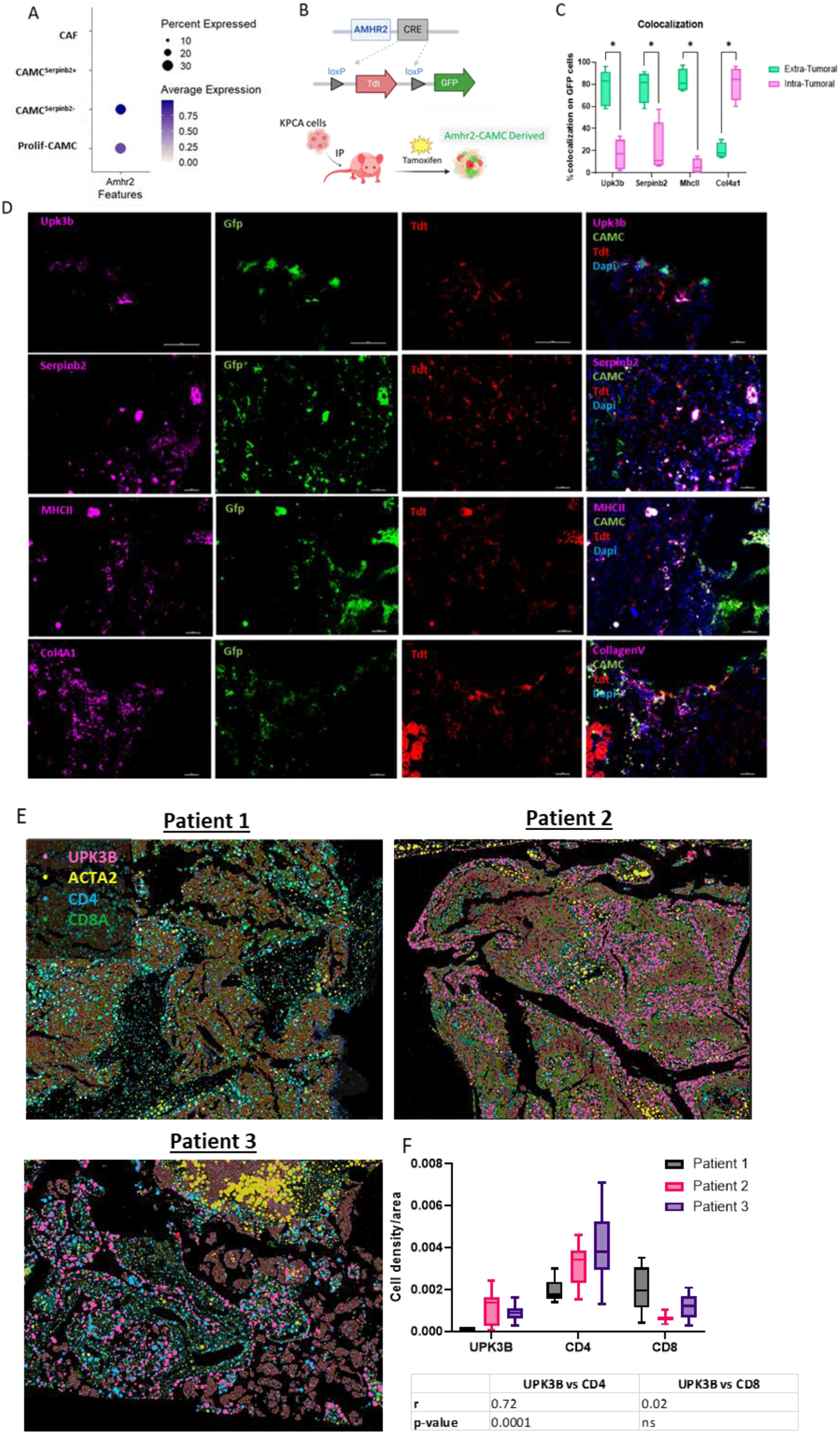
spatial distribution of CAMCs in the ovarian immune tumor microenvironment. A) Dotplot showing *Amhr2* expression in CAMC subtypes. B) Schematic of the Amhr2-lineage tracing experiment using Amhr2-CreERT2-mTmG mice. Tamoxifen was administered intraperitoneally at 20 mg/kg every other day (days 3, 5, and 7) during KPCA tumor progression to induce GFP expression in Amhr2-expressing cells. C) Quantification of immunofluorescence signal intensity for UPK3B, SERPINB3, MHC-II, and COL4A1 in both extra-tumoral (surface, 50µm thickness contour) and intra-tumoral regions of tumor sections from mTmG mice. Quantification was performed using ImageJ. N=3. D) Representative immunofluorescence images of KPCA tumor sections from Amhr2-CreERT2-mTmG mice showing lineage-traced Amhr2^+^ cells (green) and stromal cells (red). Tumor sections were stained for UPK3B, SERPINB2, MHC-II, and COL4A1 (purple) to visualize CAMC localization and marker expression within the tumor microenvironment. Scale bars: 100 µm. E) Representative spatial transcriptomics image (Xenium platform) from a high-grade serous ovarian cancer (HGSOC) patient tumor showing expression of *CD4*^+^ and *CD8*^+^ T cells, CAMC-associated transcripts (*UPK3B, SERPINB2*), and CAF markers (*ACTA2*). F) Quantification of cell density across 12 manually selected regions of interest (ROIs) spanning the tumor section. Correlation analysis was performed between *UPK3B* expression and *CD4^+^/CD8^+^* T-cell infiltration.

We have previously described AMHR2+ CAMCs located at the tumor surface in human high-grade serous ovarian tumors^13^. We sought to further refine the spatial distribution of CAMC subtypes and their proximity to immunocytes using spatial transcriptomics on HGSOC tumors from untreated patient using the Xenium platform. This analysis confirmed that UPK3B+ CAMCs were predominantly at the tumor surface and within tumor invaginations (Fig. 4E). Given their location at the tumor periphery and their MHCII expression, we investigated the spatial correlation between CAMCs and lymphocyte populations. Specifically, we correlated CAMC abundance (via UPK3B expression) with proximity to T-cell subtypes, focusing on CD4^+^ and CD8^+^ T cells. Regions of interest (ROI) with high expression of the mesothelial marker UPK3B were associated with a higher proportion of CD4^+^ T cells (r = 0.72, p < 0.0001) compared to areas lacking in CAMCs (Fig. 4F). In addition, ROI devoid of CAMCs exhibited increased CD8^+^ T cell infiltration, which could imply a more cytotoxic immune landscape in areas without CAMC influence (Fig. 3E/F). These data suggest that CAMCs may contribute to the spatial heterogeneity of the immune landscape within ovarian tumors.

### CAMC^Serpinb2+^ cells inhibit lymphocyte infiltration and promote an immunosuppressive T cell

To further investigate the effect of CAMC^Serpinb2+^ cells on immune infiltration, we used a 3D heterotypic spheroid model composed of KPCA ovarian cancer cells mixed with either immortalized (iMCs) or iMCs differentiated into CAMCs (iCAMC^Serpinb2+^) at a 1:1 ratio (2 × 10^5^ cells each). iCAMC^Serpinb2+^ were generated by stimulating iMCs during 9 days with three batches of cancer cell–conditioned media (72h each) from KPCA cells (Fig. 5A), thereby efficiently promoting their reprogramming into iCAMC^Serpinb2+^ cells (Fig. 5B). To compare the effect of iMCs and iCAMC^Serpinb2+^ on immunocyte infiltration, we performed only short-term experiments, as longer co-cultures would be expected to lead to the reprogramming of iMCs into CAMCs from contact with KPCA cells within the spheroid. Freshly isolated murine splenocytes (1 × 10^5^ cells) were added to the culture one day after spheroid formation. Spheroids were harvested after six days of culture, cryosectioned, and subjected to immunofluorescent staining (Fig. 5C). iCAMC^Serpinb2+^ labeled in red (tdTomato^+^), were distributed throughout the spheroid when co-cultured with cancer cells (Fig. 5C). Immune cells identified by immunofluorescent stain of CD45^+^ (shown in yellow), infiltrated more extensively in spheroids containing iMCs, than those containing iCAMC^Serpinb2+^, where they were largely excluded and remained confined to the periphery (Fig. 5C). Additionally, spheroids formed with iCAMC^Serpinb2+^ had a trend of larger diameter compared to those formed with iMCs (p<0.057) (Fig. 5D), suggesting the former may promote proliferation.

**Figure 5:**
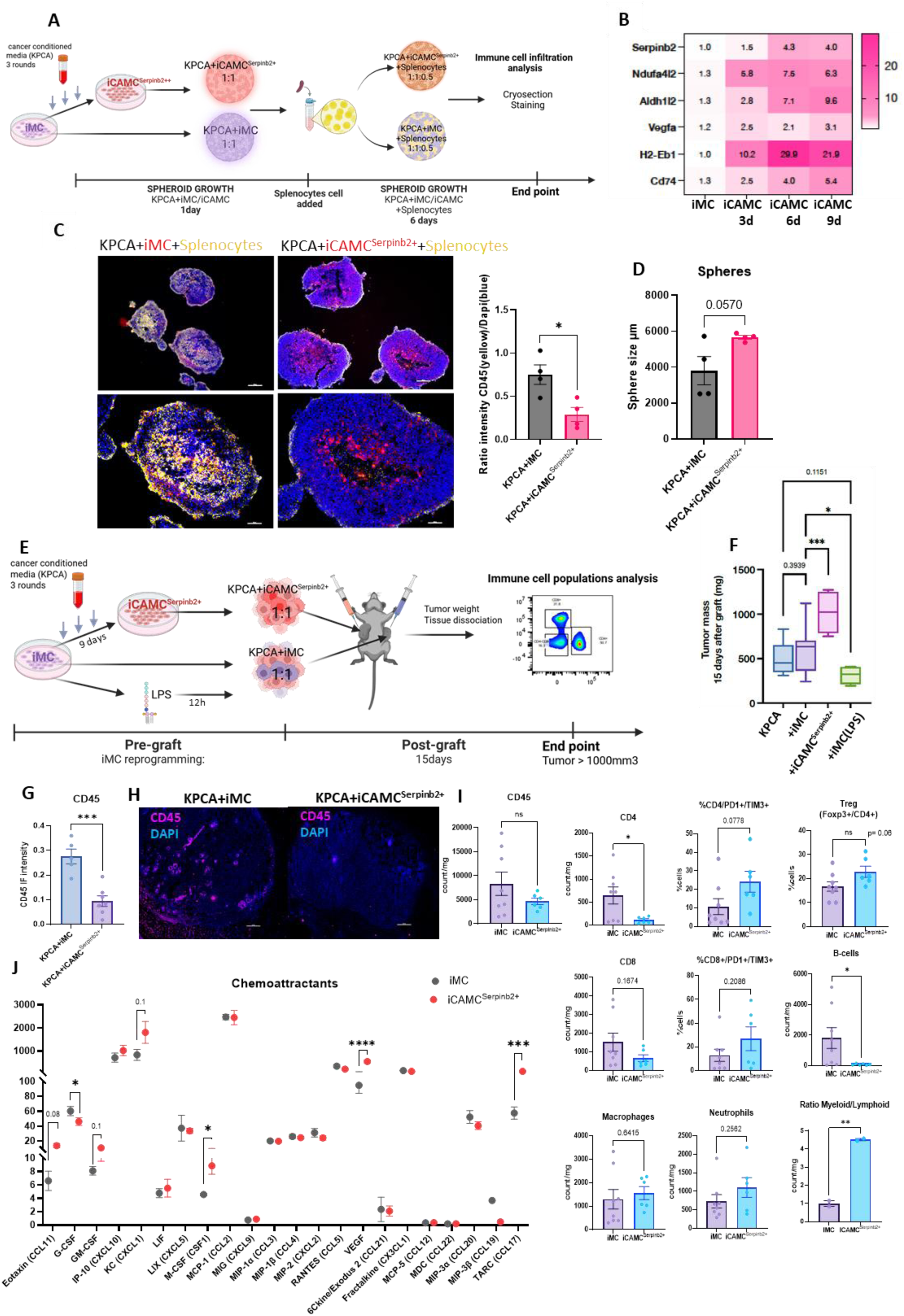
iCAMC^Serpinb2+^ enhance tumor growth and alter immune cell composition in the tumor microenvironment. A) Schematic representation of the experimental design for KPCA sphere formation in the presence of iMCs, or iCAMC **^Serpinb2+^**, followed by co-culture with splenocytes for 6 days. B) Heatmap of fold changes in CAMC^Serpinb2+^ markers and MHCII expression in mesothelial cells after 1, 2, or 3 stimulations with KPCA-conditioned media (harvest on days 3, 6, 9 respectively). C) Representative immunofluorescence images of 3D spheroids composed of KPCA and either immortalized mesothelial cells (iMCs, left panels) or immortalized cancer-associated mesothelial cells (iCAMC^Serpinb2+^, right panels) co-cultured with naives splenocytes freshly harvested form the spleen of C5J/BL6 mice. Spheres are abelled with anti-CD45 (yellow), immortalized mesothelial cells (iMC and iCAMC^Serpinb2+^) in red (tdTomato eporter) and nuclei in blue (DAPI). Scale bars = 100 µm. Bar graph quantifies the relative splenoctyes cell nfiltration (CD45 intensity) within the spheroid core (ratio of CD45^+^ intensity normalized to Dapi intensity). Data represent mean ± SEM of N = 4 spheres. Statistical significance was determined by unpaired two-tailed-test; **p < 0.01. D) Measure of the perimeter of sphere (µM), N=4. E) Schematic representation of the experimental design in which KPCA cells were grafted, mixed with iMCs, MC pre-situmulated with LPS(1ug/ml, during 12h) or with iCAMC^Serpinb2+^ (3 rounds of KPCA-conditioned media during 9 days). F) Tumor weights measured 15 days after subcutaneous grafting of KPCA cells mixed with either iMCs, MCs+LPS(1ug/ml) stimulated during 12h or iCAMC^Serpinb2+^, injected on contralateral sides of the same animal. p < 0.05, N = 4). G) Quantification of CD45^+^ immune cell infiltration based on immunofluorescence staining, comparing KPCA+iMC and KPCA+ iCAMC^Serpinb2+^ tumors. H) Representative images illustrate CD45+ staining in KPCA+iMC and KPCA+ iCAMC^Serpinb2+^ tumors. Scale bars: 100 µm. I) Immunophenotyping of tumor-infiltrating immune cells at day 15 post-grafting, as assessed by flow cytometry. Quantification includes lymphocyte populations (CD4^+^ T cells, CD8^+^ T cells, B cells), myeloid populations (macrophages, neutrophils), regulatory T cells (CD4^+^FOXP3^+^), and exhausted T cells (PD-1^+^TIM3^+^). N= 4. J) Analysis of the secretome of iMCs and iCAMC^Serpinb2+^ using multiplex ELISA (Eve Technologies), measuring he concentration of chemoattractant cytokines (pg/ml protein) in supernatants collected after 24 hours of culture. N = 3 (2 replicates).

To evaluate the iCAMC^Serpinb2+^ dependent impact on tumor microenvironment *in vivo*, we grafted KPCA cancer cells subcutaneously (2.10^6^), either alone or mixed (ratio 1:1) with iMCs, iCAMC^Serpinb2+^, or iMCs pretreated with LPS overnight (12 h) and washed three times prior to grafting (Fig. 5E). The subcutaneous model was chosen to exclude endogenous mouse peritoneal MCs, which are absent in this location. Tumors formed by KPCA cells mixed with iMCs did not show a significant increase in mass compared with KPCA cells alone, whereas co-grafting with iCAMC^Serpinb2+^ resulted in a significantly larger tumor mass (1.7-fold increase compared with iMCs; Fig. 5F).

Interestingly, iMCs pre-stimulated with LPS before grafting led to a marked reduction in tumor growth (1.9-fold decrease compared with iMCs; Fig. 5F). Together, these results indicate that cancer-derived factors reprogram mesothelial cells toward a pro-tumorigenic phenotype, while LPS treatment do not recapitulate this effect.

To better understand the influence of iCAMC^Serpinb2+^ state on the immune cell phenotype, we performed Immunostaining and compared the KPCA tumor mixed either iMC or CAMC^Serpinb2+^ (Fig. 5F) and showed a reduction of CD45+ cells infiltration in tumors with iCAMC^Serpinb2+^ (Fig. 5G/H). We next analyzed the immune subpopulations by flow cytometry in tumors derived from KPCA cells mixed with either iCAMC^Serpinb2+^ (N=4) or iMCs (N=4). Immunophenotyping of each tumor was performed using a validated panel of antibodies to identify most immune cell subtypes. The gating strategy is detailed in supplementary data (Fig. S4A/B). We found a trend for reduced total CD45+ cell count, mostly reflecting a significant reduction of CD4+ T cells and B cells, and a trend for reduced CD8+ T cells, in KPCA tumors mixed with CAMCs. Furthermore, we observed a trend of increased T regulatory (Treg) cell abundance in the iCAMC^Serpinb2+^ condition compared to iMCs. However, the myeloid infiltration was not impacted by iCAMC^Serpinb2+^, translating into a significant increase in the lymphocyte to myeloid ratio (Fig. 5I). Finally, we showed that CD4+ and CD8+ T cells exhibited an “exhausted” phenotype marked by the increase of PD1 and TIM3 markers ^32^(Fig. 5I). This was accompanied by a trend towards decreased IFNγ expression within the CD4^+^ T cell population (Fig. S4C) as well as a decrease of CD107a expression at the cell surface in CD8^+^ T cell population, indicating a lower degranulation and a potential reduction in effector and cytotoxic T cell activity (Fig. S4D) ^32^. Such a shift could reflect an immunosuppressive reprogramming of the CD4^+^ compartment, favoring tolerance over anti-tumor immunity.

To better understand how CAMCs may regulate the immune cell phenotype, we next investigated the chemokine secretion profiles of iMCs and iCAMC^Serpinb2+^. iMCs and iCAMC^Serpinb2+^ were cultured in 6-well plates at a density of 5 × 10^5^ cells per well in serum-free medium, and supernatants were collected after 24 hours to characterize their secretome using a multiplex ELISA assay (Fig. 5J). This analysis revealed a significant increase in CSF1, VEGFA, and CCL17 levels in iCAMC^Serpinb2+^ compared to iMCs. Additionally, we observed a trend toward elevated levels of GM-CSF (CSF2), CXCL1, and CCL11. Conversely, secretion of G-CSF (CSF3) was reduced in iCAMC^Serpinb2+^ (Fig. 5J). Altogether, the cytokine landscape of iCAMC^Serpinb2+^ is consistent with a tolerogenic state, and the immune exclusion of Tcells^33–35^.

### CAMC^Serpinb2+^ cells promote a tolerogenic phenotype in CD4+ T Cells

To better understand the role of CAMCs in the regulation of the immune landscape, and particularly T cells, we analyzed the communication axes across the TME using ligand–receptor inference analysis of single-cell transcriptomic data from healthy omentum and KPCA tumor (Fig. 6A). Global interaction networks revealed that in the healthy omentum, MCs exhibited no detectable interactions with T cells, suggesting a limited role in T cell modulation under non-pathogenic conditions despite their MHC-II expression (Fig. 6A). In contrast, within the tumor context, CAMCs emerged as active signaling participants, engaging specific interactions with T cells as well as cancer cells and stromal cells (Fig. 6B). The scale and specificity of these CAMC–T cell connections contrast starkly with the negligible interaction landscape observed for MCs. To evaluate the ability of iCAMC^Serpinb2+^ to modulate T cell activity compared to iMCs, we co-cultured them with naïve T cells isolated from spleen for 72hours. Expression of *Upk3b* and *Serpinb2* was evaluated in iMC and iCAMC^Serpinb2+^ by qPCR during co-culture with naïve T cells (Fig. 6C). In this co-culture setting, the presence of splenocytes further enhanced the iCAMC^Serpinb2+^ signature, marked by reduced *Upk3b* expression and increased *Serpinb2* expression at 24h, and 72h (Fig. 6C). We next evaluated T-cell markers associated with activation and regulation (CD69^+^, IL-2^+^, IFNγ^+^, GZMB^+^, FOXP3^+^) after co-culture with iMCs or iCAMC^Serpinb2+^ using flow cytometry (Fig. 6D). No differential effects were observed, suggesting that neither iMCs nor iCAMC^Serpinb2+^ can activate naïve T cells in the manner of professional antigen-presenting cells.

**Figure 6:**
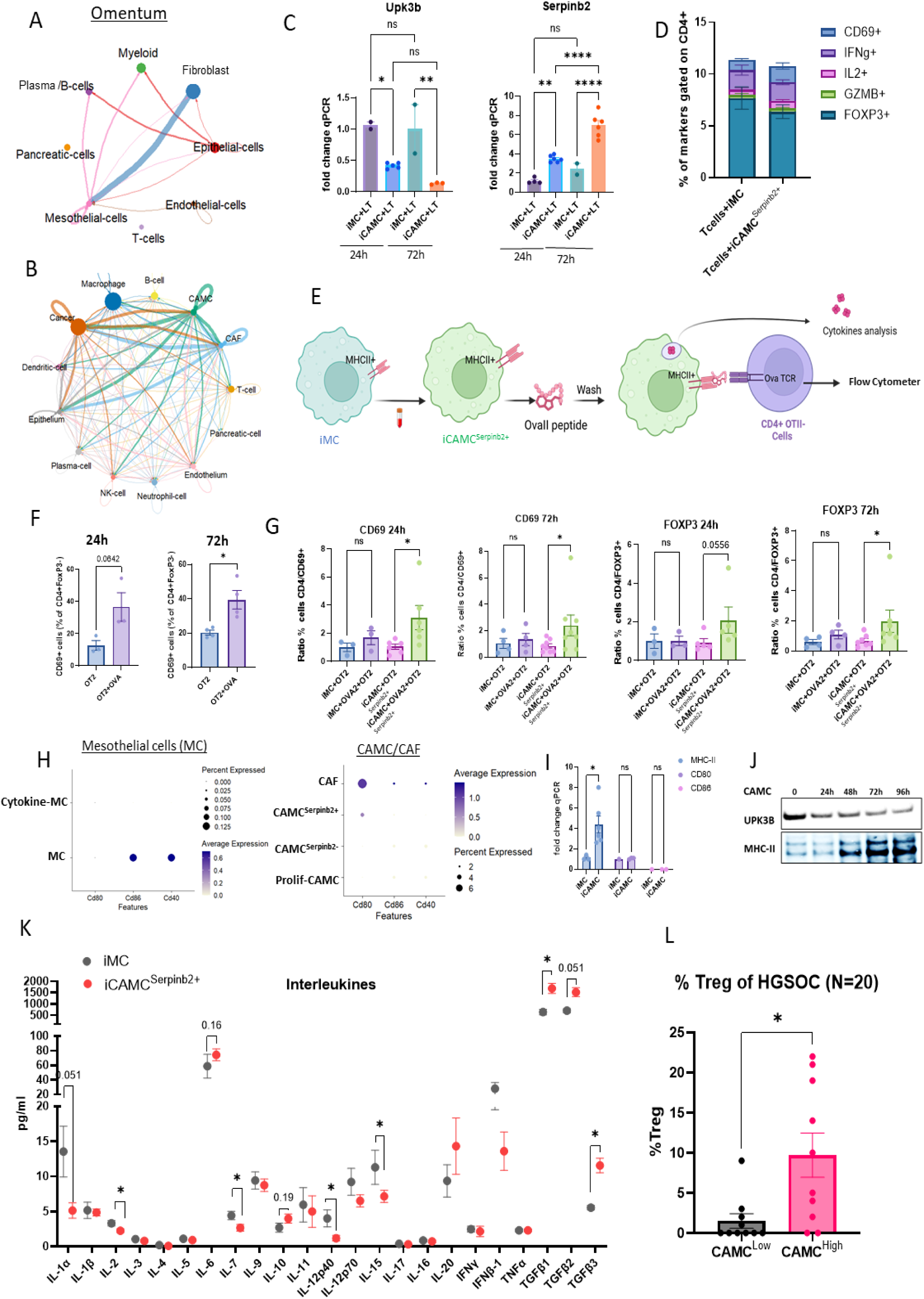
Functional characterization of mesothelial cells and their immunomodulatory interactions with T cells. A) Network representation of predicted cell-cell interactions based on ligand-receptor pair analysis on cell population from mice omentum. B) Network representation of predicted cell-cell interactions based on ligand-receptor pair analysis on KPCA tumor populations. C) qPCR analysis of *Upk3b* and *Serpinb2* expression in iMCs and iCAMC^Serpinb2+^ during co-culture with T cells at 24 and 72h. D) Flow cytometry analysis of T-cell activation and regulatory markers (CD69^+^, IL-2^+^, IFN-γ^+^, GZMB^+^, and FOXP3^+^) on CD4^+^ T cells isolated from autologous spleens and co-cultured with iMCs or iCAMC^Serpinb2+^ for 72h. E) Schematic of the experimental design for antigen-specific T-cell activation using OVA^2^-TCR transgenic CD4^+^ T cells co-cultured with iMCs or iCAMC^Serpinb2+^ in the presence or absence of OVA^2^ peptide. F) Flow cytometry analysis of CD69 expression in CD4^+^ (FOXP3-) OVA^2^-specific T cells co-cultured with iMCs or iCAMC^Serpinb2+^, normalized to OVA^2^-negative control conditions at 24h and 72h. G) Flow cytometry analysis of CD69 and FOXP3 expression in CD4^+^ OVA^2^-specific T cells co-cultured with iMCs or iCAMC^Serpinb2+^, normalized to OVA^2^-negative control conditions. H) Dot plot depicting the expression of genes encoding for co-stimulatory receptors of MHCII (*Cd80, Cd86, CD40*) based on scRNA-seq data from normal mesothelial cells (MCs) from healthy mouse omentum and CAMC subtypes in KPCA tumors. I) qPCR analysis of MHCII and co-stimulatory receptor expression in iMCs stimulated with KPCA-conditioned media. J) Western blot showing time-dependent expression of MHCII molecules in iMCs following stimulation with KPCA-conditioned media ranging from 0h to 96h. K) Secretome profiling of iMCs and iCAMC^Serpinb2+^ using multiplex ELISA (Eve Technologies) to quantify interleukin levels (pg/mg protein) in supernatants collected 24h post-seeding. Cells were plated at 5 × 10^5^ cells/well in 6-well plates. Data represent more than six independent biological replicates. L) Proportion of regulatory T cells (*CD4+FOXP3+*) in the TME of 20 HGSOC stratified by CAMC levels (CAMC^high^) or not (CAMC^low^) based on the median *UPK3B* expression.

To evaluate whether iCAMC^Serpinb2+^ can modulate CD4^+^ T-cell responses based on their increased MHCII expression following stimulation with cancer-secreted factors, we performed an antigen-specific co-culture assay. iMCs or iCAMC^Serpinb2+^ were pulsed with the Ova2 peptide, washed to remove excess of antigen, and incubated with splenocytes from OT2 mice, whose CD4^+^ T cells express a TCR specific for the Ova2 epitope (Fig. 6E)^36^. After 24h and 72h, splenocytes were harvested, and the CD4^+^ T-cell population was analyzed by flow cytometry to assess activation and polarization using an established immuno-phenotyping panel (Fig. S4). As an experimental control, splenocytes incubated with Ova peptide showed increased expression of the activation marker CD69 at both 24h and 72h (Fig. 6F). Compared to iMC, the iCAMC^Serpinb2+^ induced a marked upregulation of CD69 on CD4^+^ T cells at both time points (24h and 72h), which was correlated with increased MHCII expression (Fig 6G/H/I). In parallel, we also observed an increased proportion of CD4^+^ T cells expressing the transcription factor Foxp3, suggesting that iCAMC^Serpinb2+^ may promote the differentiation or expansion of regulatory T cells (Tregs) (Fig. 6G). We further analyzed MHCII expression along with co-stimulatory receptors (CD80/CD86 and CD40) to determine whether iCAMC^Serpinb2+^ were capable of modulating CD4^+^ T cells (Fig. 6H/I). While MHCII enables antigen presentation, co-stimulatory receptors and cytokines are required to deliver the second and third activation signals, respectively. The scRNA-seq analysis revealed MHCII expression in both iMCs and iCAMC^Serpinb2+^ (Fig. 6H), which was confirmed by qPCR and Western blot (Fig. 6I). However, barely detectable expression of co-stimulatory receptors was observed, suggesting suboptimal activation signals delivered by iCAMC^Serpinb2+^ to T cells (Fig. 6H). Together, these data suggest that CAMCs can display antigens on MHCII but likely lack the co-stimulatory signals necessary for full T cell activation.

To investigate whether CAMCs can mediate CD4^+^ T cell polarization through their cytokine secretion, we performed a cytokine array assay comparing supernatants from iMCs and iCAMC^Serpinb2+^ cultured under serum-free conditions (24-hour incubation). The multiplex analysis revealed a significant reduction of several pro-inflammatory cytokines in iCAMC^Serpinb2+^ supernatants, including IL-1α (fold change: 2.6), IL-2 (fold change: 1.5), IL-12p40 (fold change: 3.4), IL-15 (fold change: 1.6), and IL-7 (fold change: 1.7), all of which are commonly associated with promoting Th1 polarization. Conversely, IL-10, a key immunosuppressive cytokine, showed a tendency to increase in iCAMC^Serpinb2+^ supernatants (fold change: 1.4) associated with significant increase of Tgfb1 (fold change: 2.6), suggesting a shift toward a tolerogenic cytokine profile (Fig. 6K). Together with the induction of Foxp3 expression, these results support the hypothesis that iCAMC^Serpinb2+^ may contribute to immune evasion in the tumor microenvironment by skewing CD4^+^ T cell differentiation away from effector Th1 responses and toward a regulatory phenotype.

We next analyzed the proportion of Tregs (CD4+/FOXP3+) in the TME of 20 untreated HGSOC patients using the following scRNA-seq dataset (GSE147082, GSE235931, GSE241221), Patients were stratified into two groups based on TMEs enriched with all CAMCs (as defined by UPK3B+ median expression: CAMC^Low^ (N=10), CAMC^High^ (N=10)). We observed a significant enrichment of Tregs in the CAMC^High^ population (Fig. 6L), corroborating our observations of increased Treg proportions in the KPCA mouse model.

### CAMC^Serpinb2+^ Signatures Drive Therapy Resistance and Impair Immunotherapy

Based on the impact of CAMCs on reducing immune infiltration and immunosuppressive effects, we then assessed whether the CAMC Serpinb2+ subtype influenced the response to immunotherapy. First, we assessed the proportion of CAMC subtypes in KPCA tumours classified as sensitive or resistant to the triple combination of Prexasertib, anti-PD1 and anti-CTLA4 (PPC), as reported in Iyer et al., 2021, and in HGSOC patients sensitive or resistant to platinum-taxane treatment. CAMCSerpinb2+ cells were highly enriched in resistant models compared to sensitive models, both in the syngeneic mouse model and in HGSOC patients, while CAMCSerpinb2- cells were reduced. We also observed that CAFs were more abundant in resistant models (Fig. 7A/B).

**Figure 7.**
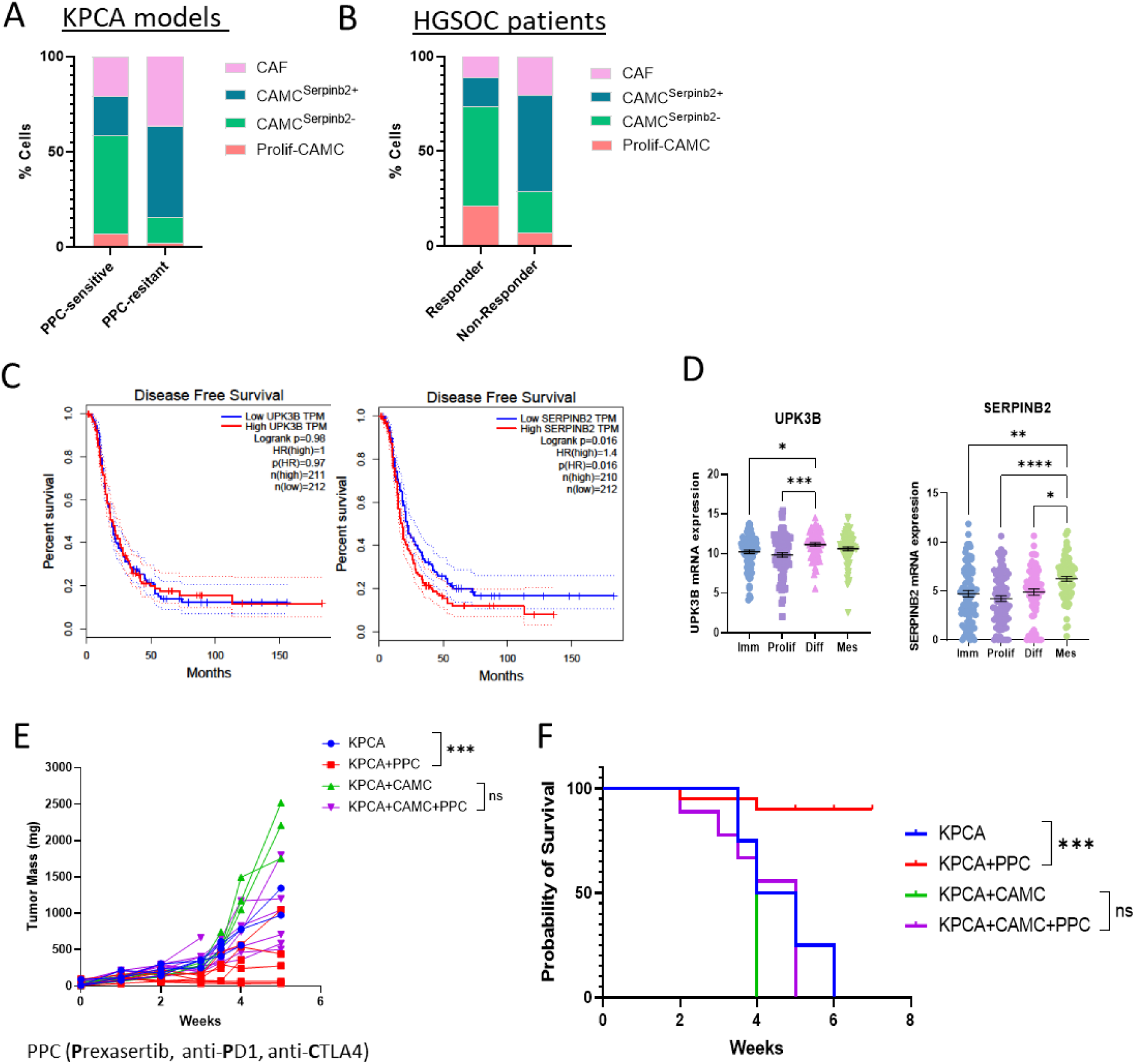
CAMC^Serpinb2+^ Cells Reduce Immunotherapy Efficacy. A) Proportion of CAMC subtypes and CAFs in KPCA tumors classified as sensitive or resistant to the triple PPC therapy (prexasertib, 10 mg/kg; anti–PD-1, 50 µg/mouse; anti–CTLA-4, 50 µg/mouse). B) Proportion of CAMC subtypes and CAFs in platinum–taxane–sensitive versus –resistant HGSOC patients from the Loret *et al.* scRNA-seq dataset (GSE201047). C) Expression of UPK3B and SERPINB2 across molecular histotypes in TCGA HGSOC tumors (N = 308). D) Kaplan–Meier survival analysis of TCGA-OV patients stratified by high versus low UPK3B and SERPINB2 expression. E) Tumor growth (mg) of subcutaneously grafted KPCA cells (4×10⁶) alone (N = 4) or mixed 1:1 with CAMCs (2×10⁶) and treated weekly with PPC (N = 8). F) Kaplan–Meier survival curves comparing PPC treatment in mice bearing KPCA tumors versus KPCA + CAMC co-grafts.

Then to assess the prognostic relevance of CAMC-associated genes, we analyzed the association between *SERPINB2* and *UPK3B* expression levels and and survival outcomes in ovarian cancer patients from The Cancer Genome Atlas (TCGA) dataset. Kaplan–Meier analysis of the TCGA OV-cohort revealed that patients with high *SERPINB2* expression had significantly worse disease-free survival compared with those with low expression (HR = 1.4, p = 0.016; Fig. 7C). This association suggests that SERPINB2 expression level may reflect the presence of abundant CAMC^Serpinb2+^ cells, which we have shown may contribute to an immunosuppressive microenvironment and are associated with the mesenchymal molecular subtype. In comparison, UPK3B had no correlation with prognosis in HGSOC (Fig. 7C).

We next sought to evaluate if *UPK3B* and *SERPINB2* overexpression were associated with a particular ovarian cancer molecular histotype^37^ (Fig. 7D). To do so we compared UPK3B and SERPINB2 transcript levels from an RNAseq dataset of the TCGA ovarian cancer UNC hub (n=308) annotated with the molecular subtypes as proliferative (Prolif), immunoreactive (Imm), differentiated (Diff), and mesenchymal (Mes), and found that *UPK3B* was significantly higher in the differentiated subtype compared to proliferative and immunoreactive (Fig. 7D). In contrast, SERPINB2 was significantly higher in the mesenchymal subtype compared to others (Fig. 6M), suggesting *SERPINB2* overexpression may be associated with mesenchymal status and worse survival outcome in human tumors^37^.

To directly assess the impact of CAMCs on therapy response, KPCA tumors were grafted subcutaneously either alone (4×10⁶ cells) or mixed 1:1 with CAMCs (2×10⁶ each) and treated weekly with the PPC combination. In KPCA-only grafts, PPC treatment significantly suppressed tumor growth, at 4 weeks reduction in tumor mass compared to untreated controls (p < 0.01; Figure 7E). In contrast, PPC treatment in KPCA + CAMC co-grafts showed no significant decrease in tumor growth, indicating that the presence of CAMCs abrogates the therapeutic effect. Consistently, PPC treatment rescued 75% of mice bearing KPCA tumors (2/8 mice died), whereas no protective effect was observed in mice with KPCA + CAMC co-grafts (all 8 mice succumbed at 6 weeks; Figure 7F). These findings demonstrate that CAMCs confer robust resistance to PPC therapy.

Collectively, these data support a model in which CAMCs actively drive immune evasion by suppressing pro-inflammatory signals while promoting tolerogenic cues. This environment may facilitate the expansion or stabilization of Tregs within the tumor microenvironment, dampening anti-tumor immunity and ultimately contributing to tumor progression and poor response to immunotherapy.

## DISCUSSION

We hypothesized that CAMCs are a functionally distinct and dynamic component of the high-grade serous ovarian cancer TME, whose differentiation states critically influence immune cell regulation and contribute to tumor immune evasion. Our objectives were to characterize the dynamics of CAMCs in the TME, and define their immunoregulatory functions and interactions with immune cell populations and ultimately their effect on immunotherapy responses.

By employing primary and immortalized syngeneic mouse HGSOC models and analyzing CAMCs in patient HGSOC tumors using scRNAseq and TCGA datasets, we identified distinct CAMC states and correlated their abundance with the immune cell repertoire within the TME.

One of the key findings of our study was the identification of a unique “metabolic” signature of CAMC observed in the KPCA tumors from a syngeneic mouse model and in human HGSOC tumors, which we defined based on SerpinB2+ expression (CAMCSerpinb^2+^). This CAMC state was characterized by a transcriptional signature marked by increased expression of metabolic and stress-response genes such as *Serpinb2*, *Ndufa4l2*, *Adm2*, *Asns*, *Aldh1l2*, *Ero1l*, *Phgdh*, and *Mthfd2*. Although the CAMC^Serpinb2+^ cluster is distinct in its molecular profiles, we propose that it represents a transient state of differentiation of CAMCs, rather than a static subpopulation, potentially indicating the progression of CAMC^Serpinb2-^ cells towards a CAF-like phenotype. Supporting this, conclusion *in vitro* experiments showed that MCs can acquire CAMC^Serpinb2+^ signature upon stimulation with cancer-conditioned media in a time-dependent manner. The temporal changes in gene expression likely also coincide with spatial changes in the localization of these cells. Lineage tracing of CAMCs confirmed the localization of UPK3B, SERPINB2, and MHCII at the tumor periphery. As CAMCs migrated and infiltrated the tumor core, they progressively lost expression of these CAMC-specific markers. This phenotypic shift highlights CAMC plasticity and transition toward a mesenchymal-like state, as evidenced by the upregulation of COL4A1 and ACTA2, markers of CAF subtypes. Such a transition is consistent with the well-documented mesothelial-to-mesenchymal transition (MMT), whereby mesothelial cells differentiate into fibroblast-like cells in both fibrosis and peritoneal metastasis contexts^10,24,38–41^. This mesothelial plasticity could explain the emergence of me

Importantly, the CAMC^Serpinb2+^ gene expression signature is tumor-specific and dependent on cancer-secreted factors. Its absence in normal mesothelial cells or upon LPS stimulation indicates that this program is not a general stress or inflammatory response^31,42^, but a state actively induced by cancer cells. This specificity suggests that CAMC^Serpinb2+^ cells represent a tumor-adapted pathogenic mesothelial cell state.

We next characterized the role of CAMCs in shaping the tumor immune landscape. We hypothesized that CAMC^Serpinb2+^ secretome may influence the immune repertoire of the tumor. CAMC^Serpinb2+^ secrete elevated levels of M-CSF (CSF1) and G-CSF (CSF3), two cytokines that promote the recruitment and functional polarization of tumor-associated macrophages and neutrophils, respectively, both major immunosuppressive populations in ovarian cancer ^43,44^. Moreover, CAMCs produced high levels of CCL17, a CCR4 ligand that selectively attracts Tregs, reinforcing a regulatory niche within the TME^45^. In line with this, analysis of HGSOC patient samples revealed that tumors with higher CAMC^Serpinb2+^ abundance exhibited significantly increased Treg infiltration, a feature associated with poor prognosis. In addition to these chemokines and growth factors, CAMC^Serpinb2+^ showed a significant reduction in IL-2, IL-15, and IL-12p40, cytokines normally required for effector T cell proliferation, survival, and Th1 polarization, while concomitantly increasing IL-10 and TGFβ1, two potent immunosuppressive cytokines. Taken together, this biased secretory repertoire positions CAMC^Serpinb2+^ as central architects of an immune-suppressive, tumor-permissive microenvironment by promoting Treg recruitment, impairing effector T cell function, and excluding adaptive immune cells from tumor sites. Strikingly, VEGF was also enriched in CAMC^Serpinb2+^ conditioned media, providing a dual mechanism of action by driving angiogenesis while simultaneously restricting effector T cell infiltration, thereby contributing to immune exclusion^34^.

Beyond their secretory activity, CAMC^Serpinb2+^ exert a direct influence on CD4^+^ T cell responses through MHCII antigen presentation. Although CAMCs express MHCII in a time-dependent fashion, they lack canonical co-stimulatory molecules such as CD80, CD86, and CD40, positioning them as non-professional antigen-presenting cells. In functional assays, CAMCs were able to present antigen and induce CD69+ marker on CD4^+^ T cells, but this was accompanied by an increased frequency of FOXP3^+^ T cells, indicating a tolerogenic state. These results suggest that CAMC^Serpinb2+^ may act as an atypical antigen-presenting cell that divert CD4^+^ T cell responses toward anergy or regulatory fates, thereby dampening anti-tumor immunity. This parallels findings from pancreatic cancer, where antigen-presenting CAFs have been shown to promote T cell tolerance^46,47^.

Spatial transcriptomic analysis of HGSOC tumors from four patients revealed that regions enriched in surface CAMCs (*UPK3B*^+^) strongly correlated with CD4^+^ T cells but appeared to restrict their infiltration into the tumor core. In contrast, regions lacking UPK3B expression exhibited higher CD8^+^ T cell infiltration, suggesting that CAMC presence influences both the composition and localization of tumor-infiltrating lymphocytes. Extending these findings, we observed that tumors enriched in CAMCs displayed reduced infiltration of CD4^+^ T cells and B cells, while favoring the accumulation of regulatory T cells (Tregs). Together, these data establish CAMCs as active contributors to the remodeling of the tumor immune microenvironment (TME), whereby their acquisition of a distinct immunomodulatory phenotype, compared with normal mesothelial cells, may support tumor immune evasion.

This study suggests that acquiring a CAMC^Serpinb2+^ signature, may correlate with both immune tolerance and increased tumor growth. Indeed, co-graqfting CAMC^Serpinb2+^ along with cancer cells increased tumor burden by 1.6-fold compared to normal MCs. In addition, our study identifies CAMC^Serpinb2+^ cells as critical mediators of resistance, both in patient tumors and in syngeneic KPCA models, CAMC^Serpinb2+^ cells were enriched in non-responder models, while CAMC^Serpinb2-^ populations were reduced. Furthermore, the CAMC^Serpinb2+^ signature was also found in the tumor microenvironment of humans HGSOC tumor and associated with worse disease-free survival, and the mesenchymal molecular phenotype associated with the worst outcome. Collectively, these findings indicate that the poor efficacy of immunotherapy is, at least in part, driven by the immunosuppressive activity of CAMCs, which identifies them as potential contributors to resistance and as therapeutic targets for restoring responsiveness to immunotherapy.

A key question remains regarding how cancer cells reprogram normal mesothelial cells into CAMCs and ultimately into CAFs, and whether these transitions are reversible. Targeting these processes could reduce tumor growth and limit immune escape. Direct elimination of CAMC^Serpinb2+^, for example, through ADCs against CAMC^Serpinb2+^ specific surface markers, may dismantle pro-tumoral niches while sparing normal tissue. Alternatively, harnessing the antigen-presenting function of CAMCs to deliver costimulatory signals could reprogram T cells toward anti-tumor responses. Defining how CAMC - T cell interactions shape immunity may open new therapeutic avenues.

In conclusion, CAMCs are emerging as crucial players in shaping the ovarian cancer microenvironment, promoting immune evasion, stromal remodeling, and ultimately contributing to disease progression. Therapeutic strategies aimed at preventing CAMC differentiation or restoring their normal functions could provide novel avenues for improving patient outcomes in HGSOC

## ACKNOWLEDGEMENT

We thank Caroline Coletti for her help in maintaining IRB and IACUC protocols and general support of research activities. We thank Patricia K. Donahoe for her general support of the laboratory.

## DATA ACCESS STATEMENT

The datasets generated and/or analyzed during the current study are available from the corresponding author on reasonable request. RNA sequencing data supporting the findings of this study are included in the article and its supplementary information files.

## AUTHORS CONTRIBUTION

Conceptualization: M.C. and D.P.; methodology: M.C., J.R-P., E.T., L.G., MCM.; investigation, M.C., D.P., J.R-P., E.T., R.M., M.-C.M.; writing original draft: M.C, D.P.; writing review & editing: N/A; funding acquisition: D.P. M.C; Resources: D.P., M.C, HA.M, V.L., L.G., N.B., PEC; supervision: M.C., D.P.

## METHODES

### EXPERIMENTAL MODEL AND SUBJECT DETAILS

#### Cell models

The KPCA cell lines were developed and described in Iyer et al.^22^ and the OV90 cells lines of malignant papillary serous adenocarcinoma were purchased from ATCC. These cells were cultured in DMEM medium supplemented with 1% of FBS and 1% penicillin and streptomycin and incubated at 37°C/5%CO2. The primary human ovarian cancer cell (ptD) was collected from ascites of patient after informed consent under Institutional Review Board-approved protocols (2007P001918/Massachusetts General Hospital) as part of a previous study^30^. The adherent cells were cultivated in DMEM:F12, supplemented with 5% of FBS and 1% of penicillin/streptomycin. The Met5a cell line was purchased from ATCC and cultured in DMEM medium supplemented with 5% FBS. Primary Mesothelial cells (MCs) were obtained from the omenta of female mice by mechanically disrupting small pieces and performing enzymatic digestion using collagenase at 37°C for 45 minutes. The pieces of tissue were placed in a collector tube in a gentlemacs dissociator (Miltenyi) for 1min of processing. The resulting cell suspension was filtered through a 70 μm filter and washed 2 times with PBS (Phosphate Buffer Saline). The adherent cells were cultured in DMEM:F12 medium, supplemented with 5% of FBS (Fetal Bovine Serum) and 1% of penicillin/streptomycin.

#### Generation of immortalized Mesothelial Cells (iMCs)

Immortalized mesothelial cells (iMCs) were generated by transducing primary mesothelial cells with a lentiviral vector encoding the SV40 large T antigen. After 72h, cells were cultured in the presence of puromycin (2 µg/mL) for two weeks to select a homogeneous, stably transduced population. The resulting iMCs were maintained in DMEM/F12 medium supplemented with 5% fetal bovine serum (FBS) and 1% penicillin/streptomycin.

#### Generation of CAMC by conditioned media treatment

Cancer-associated mesothelial cells (CAMCs) and their immortalized counterparts (iCAMCs) were generated by stimulating primary mesothelial cells or iMCs, respectively, with standardized KPCA cancer cell-conditioned medium. This medium was prepared by culturing KPCA cells to 80% confluence, replacing the medium with DMEM/F12 containing 1% FBS, and collecting the supernatant after 24 hours. The conditioned medium was centrifuged to remove cellular debris and applied to MC or iMC cultures for a minimum of 72 hours to induce a CAMC^Serpinb2+^ phenotype. For *in vivo* experiments, iCAMCs were subjected to three consecutive rounds of KPCA-conditioned media stimulation prior to the subcutaneous implantation.

#### Mouse models

This study was performed according to experimental protocols 2009N000033 and 2024000039, approved by the Massachusetts General Hospital Institutional Animal Care and Use Committee with C57BL6J mice purchased from the Jackson Laboratory. All experiments were made with 12-16 weeks female C57BL6J mice. In each experiment, we used 4 to 5 mice per group.

*<u>Amhr2-CRE-ERT2 x mTmG</u>* mice were developed in collaboration with the Jackson Laboratory (Strain #:037056 RRID:IMSR_JAX:037056). Briefly, they are the result of a knock-in of a CRE-ER fusion transgene in frame with the Amhr2 gene using a T2 exon skipping mechanism to ensure Amhr2 expression is maintained. These mice were then crossed with the Rosa26 CRE-inducible mTmG reporter as previously described^13,48,49^. Ear genotyping of Amhr2-ERT2 and mTmG reporter mice was done using the REDExtract-N-Amp tissue PCR kit (Sigma, #SLBT8193) with sets of primers design by IDT and Invitrogen (Key resource table). Tamoxifen (4-OHT, Sigma-Aldrich) treatment was used to induce CRE-ER nuclear translocation, at a final concentration of 2µM *in vitro* in female mesothelial cells isolated from omenta as described above. Only female mice carrying the Amhr2-CRE-ERT2 x mTmG were used in this study.

#### Human samples

Spatial transcriptomics analysis from tumor sections were donated by the Biological Resources Center of Montpellier Cancer Institute (ICM), France, and collected following the French regulations under the supervision of an investigator. The collection was declared to the French Ministry of Higher Education and Research (declaration number DC-2008–695). The tumor sections originate from HGSOC chemonaïve patients.

#### Cancer cell graft

Female mice were injected subcutaneously with 4 × 10⁶ KPCA cells, either alone or in combination with iMCs, iMC+LPS (1ug/ml) or iCAMC^Serpinb2+^ at a 1:1 ratio. Mice were sacrificed 17 days post-injection, or when tumor volume exceeded 1500 mm³, to evaluate tumor burden.

Prexasertib, anti-PD1, anti-CTLA4 treatment combination: Treatments began when subcutaneous tumor rich 100mm3 and were administered intraperitoneally once a week with the indicated doses during 6 weeks. For immunotherapy, anti-CTLA4 (50 μg/mouse; Bio X Cell, BE0164) and anti-PD1 (50 μg/mouse; Bio X Cell, BP0273) antibodies were used. Prexasertib (Selleckchem, S7178; 10 mg/kg) was administered in combination with the immunotherapies and was resuspended according to the manufacturer’s instructions. Control mice were injected with 10% DMSO (vehicle) and isotype antibodies.

#### Bulk-RNAseq

RNA was extracted from primary MCs and CAMCs using the Qiagen RNA extraction kit according to the manufacturer’s instructions. Library construction, sequencing, and analysis was performed by Novogene Corporation Inc. Genes with adjusted p-value < 0.05 and |log2(FoldChange)| > 1 were considered as differentially expressed. Genes listed in Table 1: RNA sequencing performed on primary mesothelial cells (MCs) and MCs stimulated with KPCA media from two clones (KPCA.B and KPCA.C) during 72h.

#### Immunofluorescence

The immunofluorescence staining was performed on fixed cells permeabilized with triton X-100 (0.1%). The cells were incubated for 1 hour in blocking buffer at room temperature with PBS-BSA (2%), followed by three washes with PBS-Tween (0.01%). Then, the cells were incubated with the primary antibody overnight at 4°C, with the antibody concentration as recommended by the manufacturer. The cells were washed and incubated for 1 hour with the specific secondary antibody. To visualize the stained proteins, the cells were counterstained with the DAPI nuclear stain 5 minutes and a final wash step with PBS was performed. The coverslips were mounted onto glass slides using Vectashield mounting medium to preserve the fluorescent signal.

#### qPCR

Total RNA was extracted from mesothelial cells using the Qiagen RNA extraction kit, and cDNA was synthesized from total RNA using SuperScript III First-Strand Synthesis System for RT-PCR (Invitrogen). Expression levels relative to the geometric mean housekeeping genes were calculated by using cycle threshold (Ct) values logarithmically transformed using the 2^−ΔCt^ function.

#### Western blot

MCs or iMCs were stimulated with cancer-conditioned media collected from KPCA cultures. Cells were incubated and sampled at different time points (0, 24, 48 and 72h). After stimulation, cells were washed with cold PBS and lysed in RIPA buffer supplemented with protease and phosphatase inhibitors. Protein concentrations were determined using a BCA assay. Equal amounts of protein (20–30 µg) were resolved by SDS–PAGE and transferred onto PVDF membranes (Millipore). Membranes were blocked in 5% non-fat dry milk in TBST for 1 h at room temperature and incubated overnight at 4 °C with primary antibodies against (specifiied target proteins, e.g., UPK3B, MHCII, SERPINB2 and B-ACTIN as loading control). After washing, membranes were incubated with HRP-conjugated secondary antibodies for 1 h at room temperature. Protein bands were visualized using enhanced chemiluminescence (ECL, Thermo Fisher Scientific).

#### Cytokine array

The cytokine profile was quantified in supernatants collected from *in vitro* cultures of iMCs and iCAMCs. Cells were seeded in 6-well plates and cultured for 24 h under serum free. Supernatants were harvested, centrifuged at 300 × g for 5 min to remove debris, and stored at −80 °C until analysis. Cytokine concentrations were measured using a multiplex ELISA panel (Eve Technologies, Calgary, Canada) according to the manufacturer’s instructions. Samples were run in triplicate in three independent experiments.

#### Spheroid growth

Spheroids were generated by seeding KPCA cancer cells (2.10^5^) either alone or mixed at a 1:1 ratio with mesothelial cells MCs or CAMCs in ultra-low attachment 96-well plates (Corning). To investigate the contribution of mesothelial cells within the tumor microenvironment (TME), spheroids of KPCA cancer cells were first cultured and subsequently supplemented with MCs (1.10^5^ cells) for 1 or 2 weeks. At the indicated time points, spheroids were collected, embedded in OCT compound, cryosectionned, and subjected to immunofluorescence staining to evaluate mesothelial cell persistence and immune cell infiltration.

To assess immune cell infiltration, KPCA cells were co-cultured with either iMCs or iCAMC^Serpinb2+^ at a 1:1 ratio in ultra-low attachment plates. Spheres were supplemented with 1 × 10^5^ freshly isolated splenocytes in complete RPMI medium and maintained for 6 days at 37 °C with 5% CO^2^. At the endpoint, spheres were harvested, cryo-embedded in OCT, sectioned at 8 µm, and subjected to immunofluorescence staining with anti-CD45 antibodies to evaluate immune cell infiltration.

#### Flow cytometry

KPCA tumors (N = 4) were dissociated into single-cell suspensions, days 15 post graft. A total of 1.10^5^ cells per well were plated in 96-well V-bottom plates. Cells were first immunostained for extracellular markers, then permeabilized using Nuclear Foxp3 Perm Buffer (according to the Biolegend’s instructions) for intracellular staining. After permeabilization, cells were washed and stained with antibodies targeting intracellular markers as recommended by the manufacturer, followed by three additional washes. Cells were then fixed with 1% paraformaldehyde. To discriminate live and dead cells, the Live/Dead reagent (Viakrom808) was used. Immunostained cells were processed on a Cytoflex flow cytometer (BD), and data analysis was performed using FlowJo software (v.10.)

#### OVA/OT-II T Cell Co-culture

iMCs or iCAMC^Serpinb2+^ were treated with 100 µg/mL Ovalbumin II (OVA II) peptide for 1 hour, washed three times, and then co-cultured at a 1:3 ratio with splenocytes from OT-II transgenic mice, which express the mouse α- and β-chains of the T cell receptor specific for OVA II and pair with the CD4 co-receptor. Co-cultures were performed in low-attachment 96-well plates for 24 or 72 hours. At the end of the culture period, cells were harvested, washed, and stained for extracellular and intracellular markers. Live/dead discrimination was performed using the Viakrom reagent. Flow cytometry acquisition was performed on a Cytoflex (BD), and data were analyzed using FlowJo software.

### QUANTIFICATION AND STATISTICAL ANALYSIS

#### Statistical analysis software

Statistical analysis was performed using GraphPad Prism 10.6 software (GraphPad Software, Inc. RRID:SCR_002798), using an independent sample t-test unless otherwise indicated or ANOVA test. P≤0.05 was considered as significant. Data are represented as mean ± standard error of the mean (SEM).

#### Image analysis

ImageJ software was used to quantify signal intensity following the pipeline described below. Each individual fluorescence channel corresponding to the proteins of interest (UPK3B, MHCII, SERPINB2, COL4A1, or ACTA2) was quantified and normalized to the DAPI channel. Tumor tissue was manually delineated, and a peripheral region extending 50 µm from the tumor contour was defined. The mean fluorescence intensity of each marker was then quantified within the selected regions.

#### Spatial transcriptomic analysis

Gene transcripts were conducted using the Xenium platform (10x Genomics) on three untreated high-grade serous ovarian cancer tumor samples. The expression levels of *UPK3B*, *SERPINB2*, *ACTA2*, *CD4*, and *CD8* transcripts were quantified across >10 representative regions of interest (ROI) per tumor to capture the spatial heterogeneity. The data was processed and analyzed using Xenium software (10x Genomics), enabling precise transcript quantification and spatial mapping of gene expression patterns. The r correlation was performed between *UPK3B* and *CD4* or *CD8* transcripts.

#### Co-immunofluorescence image analysis

Quantification of colocalization was performed using the Coloc2 plugin in ImageJ and corresponds to the percentage of area overlap between purple staining (UPK3B or MHCII or SERPINB2 or COL4A1) or and GFP staining.

#### Single-cell RNA and TCGA analysis

Transcriptomic data were extracted from the following datasets: human HGSOC (GSE235931, GSE14082, GSE241221), KPCA model (GSE233423), and healthy murine omentum (GSE134355), and analyzed in RStudio using the Seurat package. Patients were stratified into two groups based on *UPK3B* expression: CAMC-High (average expression above the population median) and CAMC-Low (average expression below the population median). The tumor microenvironments of CAMC-High and CAMC-Low patients were compared using specific markers indicative of cell phenotypes of interest, including *CD4*+ *FOXP3*+ cells. To further assess differences between groups, the percentage of cells expressing these markers was calculated using the WhichCells command in Seurat.

To assess UPK3B and SERPINB2 expression across molecular histotypes of HGSOC, we analyzed transcript levels from the TCGA ovarian cancer (OV) UNC hub RNA-seq dataset (n = 308), which includes metadata categorizing samples into proliferative, immunoreactive, differentiated, and mesenchymal subtypes. Level 3 data were obtained from the UCSC Xena repository. Transcript counts generated by Illumina HiSeqv2 were log-transformed using the formula log^2^(norm_count + 1) for downstream analysis.

#### Survival Analysis

Disease-free survival (DFS) analyses for UPK3B and SERPINB2 were performed using the GEPIA3 (Gene Expression Profiling Interactive Analysis, version 3) online tool (https://gepia3.bioinfoliu.com). This platform integrates RNA-sequencing expression data from The Cancer Genome Atlas (TCGA) and the Genotype-Tissue Expression (GTEx) projects.

For each gene of interest, patients were stratified into high- and low-expression groups according to the median expression value. Kaplan–Meier survival curves were generated to evaluate differences in DFS between groups. The log-rank test (Mantel–Cox) was used to determine statistical significance, and hazard ratios (HR) with 95% confidence intervals (CI) were reported. Graphical outputs were exported directly from the GEPIA3 interface.

#### Declaration of generative AI and AI-assisted technologies in the manuscript preparation process

During the preparation of this work the author(s) used ChatGPT for proof reading only. After using this tool/service, the author(s) reviewed and edited the content as needed and take(s) full responsibility for the content of the published article.

## SUPPLEMENTAL FIGURES

**Figure S1:**
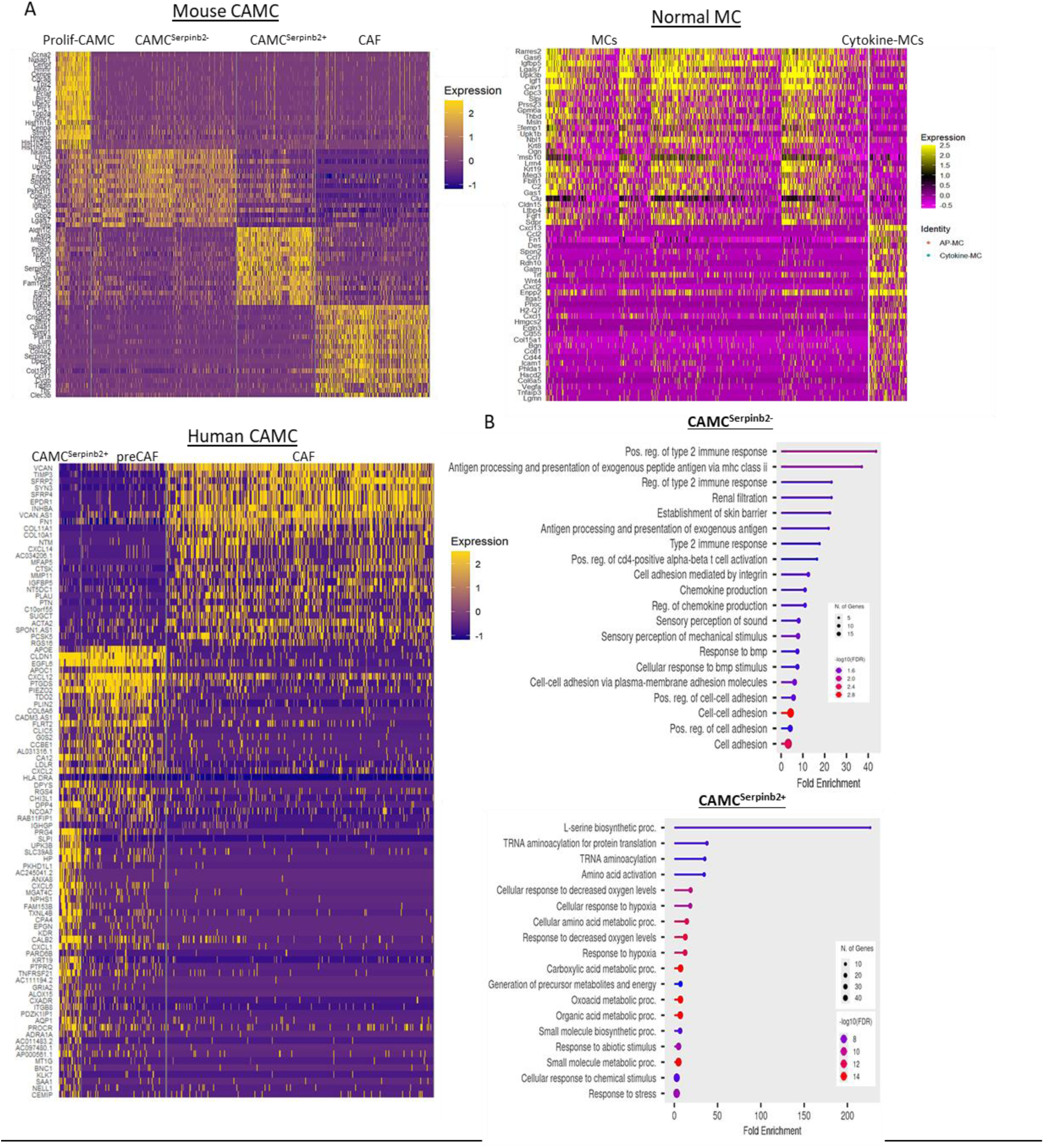
Identification of CAMC subtypes gene signature in ovarian cancer microenvironment. A) Heatmap illustrating the expression profiles of genes across mouse CAMC subtypes (Prolif-CAMC, CAMC^Serpinb2-^, CAMC^Serpinb2+^, and CAF), mouse Mesothelial cells (MC and Cytokine-MC) and Human CAMC (CAMC^Serpinb2+^, PreCAF, and CAF). Each row represents a gene expression (Log2Fc>1), while each column corresponds to individual cell, with the color intensity indicating the level of expression (yellow: high expression, purple: low expression). B) Gene Ontology (GO) enrichment analysis of differentially expressed genes (Log^2^FC > 0.5) of CAMC^Serpinb2-^ and CAMC^Serpinb2+^ from the scRNAseq of mouse CAMC clusters.

**Figure S2:**
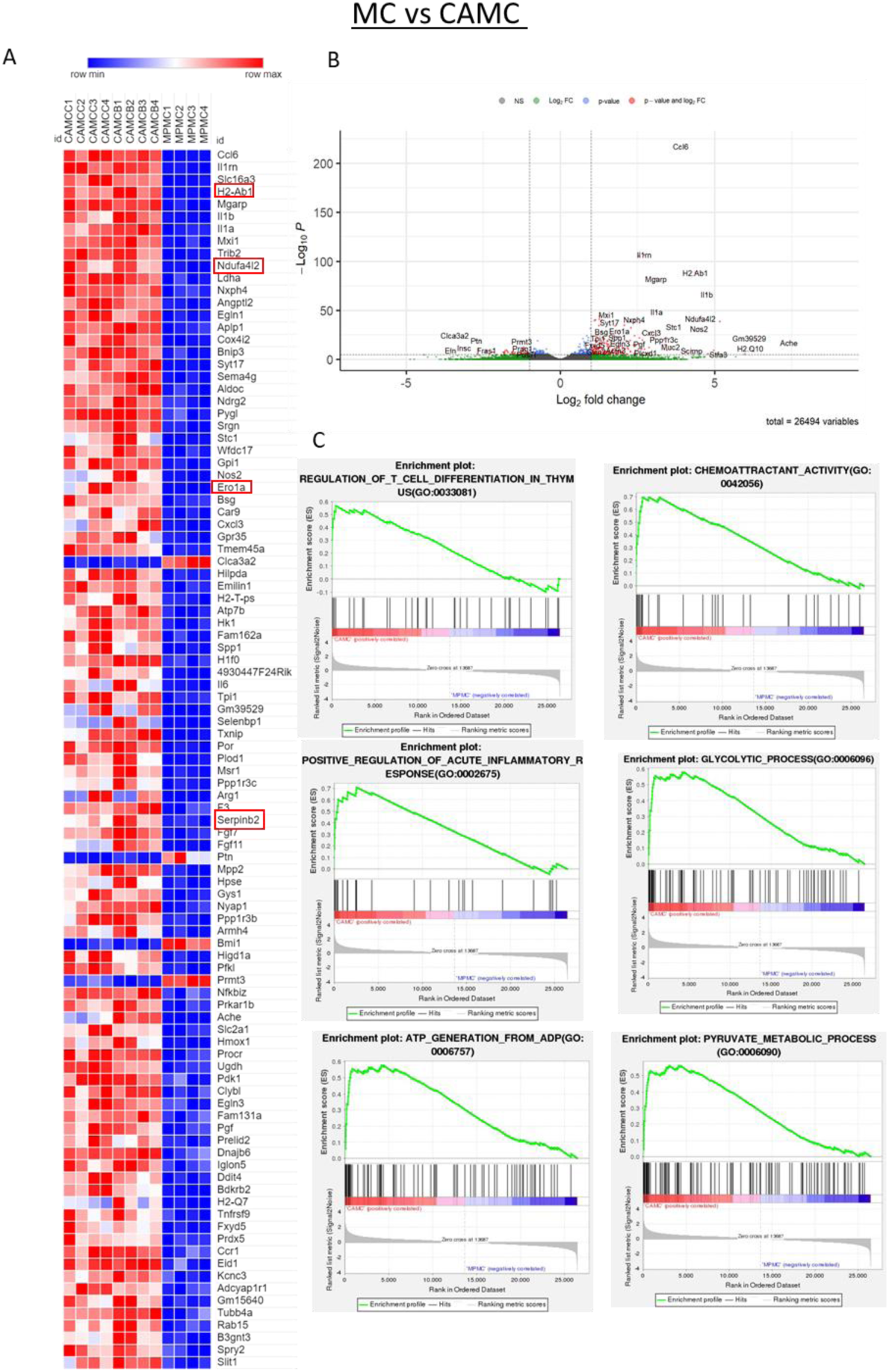
A) Heatmap showing the differential expression of genes comparing mouse primary mesothelial cells (MCs) and cancer-associated mesothelial cells (CAMCs) bulk RNA sequencing following 72h treatment of MCs with cancer-cell conditioned media (from KPCA) or vehicle control. N=2 independent experiments realized in duplicate. Differentially expressed markers between two distinct conditions, with each column representing an individual sample. The color scale indicates relative gene expression levels, ranging from high (red) to low (blue). B) Volcano plot displaying 26,494 genes expression levels in CAMCs versus MCs. Genes significantly dysregulated by log2fold change and p-value are marked in red for the upregulated ones and blue for the downregulated ones respectively. Green dots represent genes with a significant fold change but not p-value, while gray indicates non-significant changes. Dashed lines represent thresholds for significance and fold change cutoffs. C) Enrichment plots illustrating the gene set enrichment analysis (GSEA). The green curve represents the enrichment score (ES), indicating the cumulative distribution of gene hits across the ranked dataset. Genes correlated with CAMC are positively enriched (highlighted in red), while those correlated with MCs are negatively enriched (highlighted in blue).

**Figure S3:**
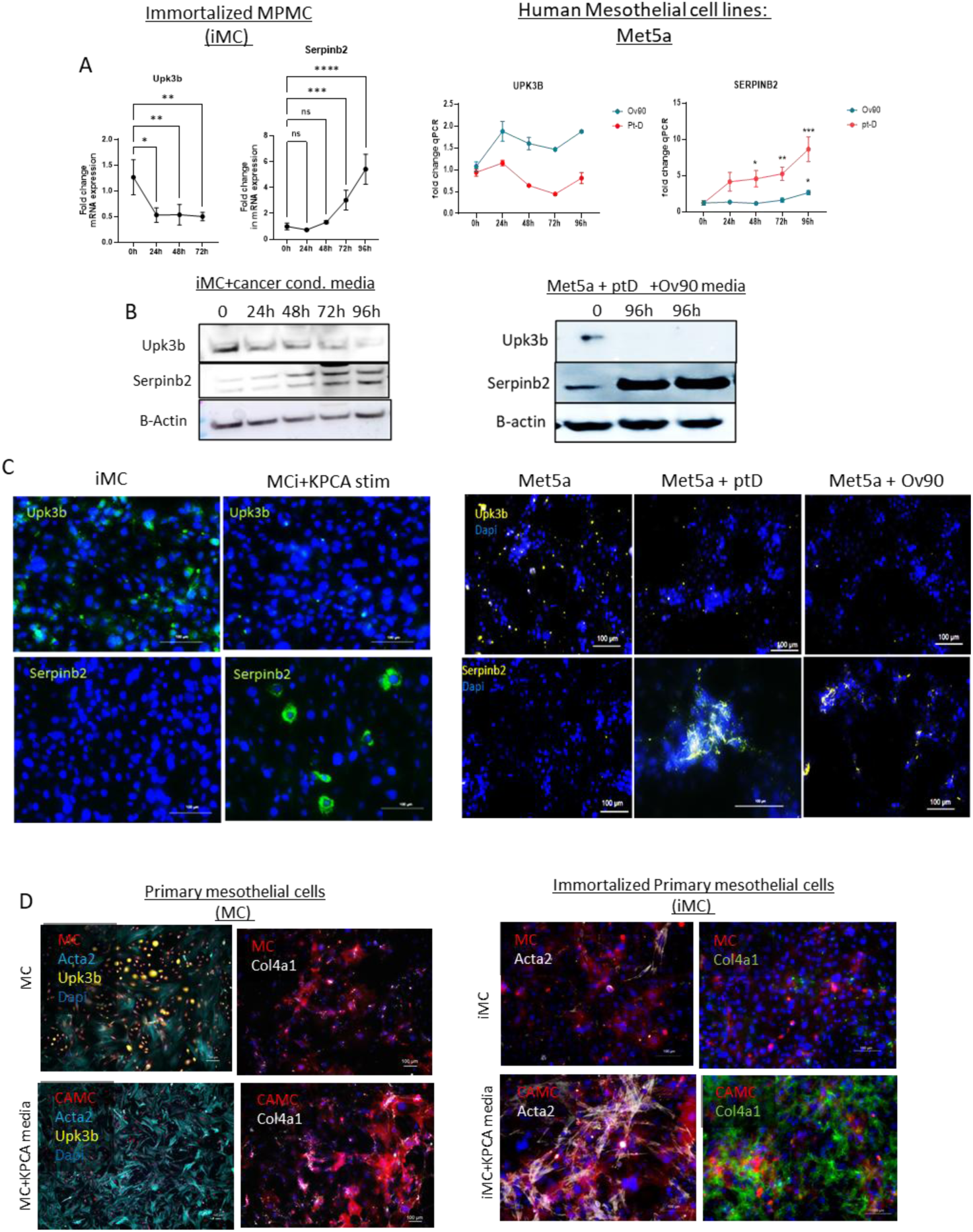
Differentiation of Mesothelial cells into CAMCs. A) Quantitative PCR (qPCR) analysis of *Upk3b* and *Serpinb2* expression in immortalized mesothelial cells (iMC) stimulated with KPCA-conditioned media and Met5a stimulated with patient D HGSOC media (ptD) or Ov90 ovarian cancer cell line in a time-dependent manner (0-96hrs). B) Western blot analysis of UPK3B and SERPINB2 expression in iMC treated with KPCA-condioned culture medium at 0h, 24h, 48h, 72h and 96h (left panel) and Met5A stimulated with conditioned media from ptD/Ov90 cell-lines versus unstimulated controls at 72h (right panel). C) Immunofluorescence analysis of UPK3B and SERPINB2 in iMC and Met5A comparing conditions stimulated with either KPCA-conditioned media or ptD/Ov90 media versus unstimulated controls at 72 h. D) Immunofluorescence staining for ACTA2 and COL4A1 in both primary MCs and immortalized MCs (iMCs) after 96 hours of exposure to KPCA-conditioned media

**Figure S4:**
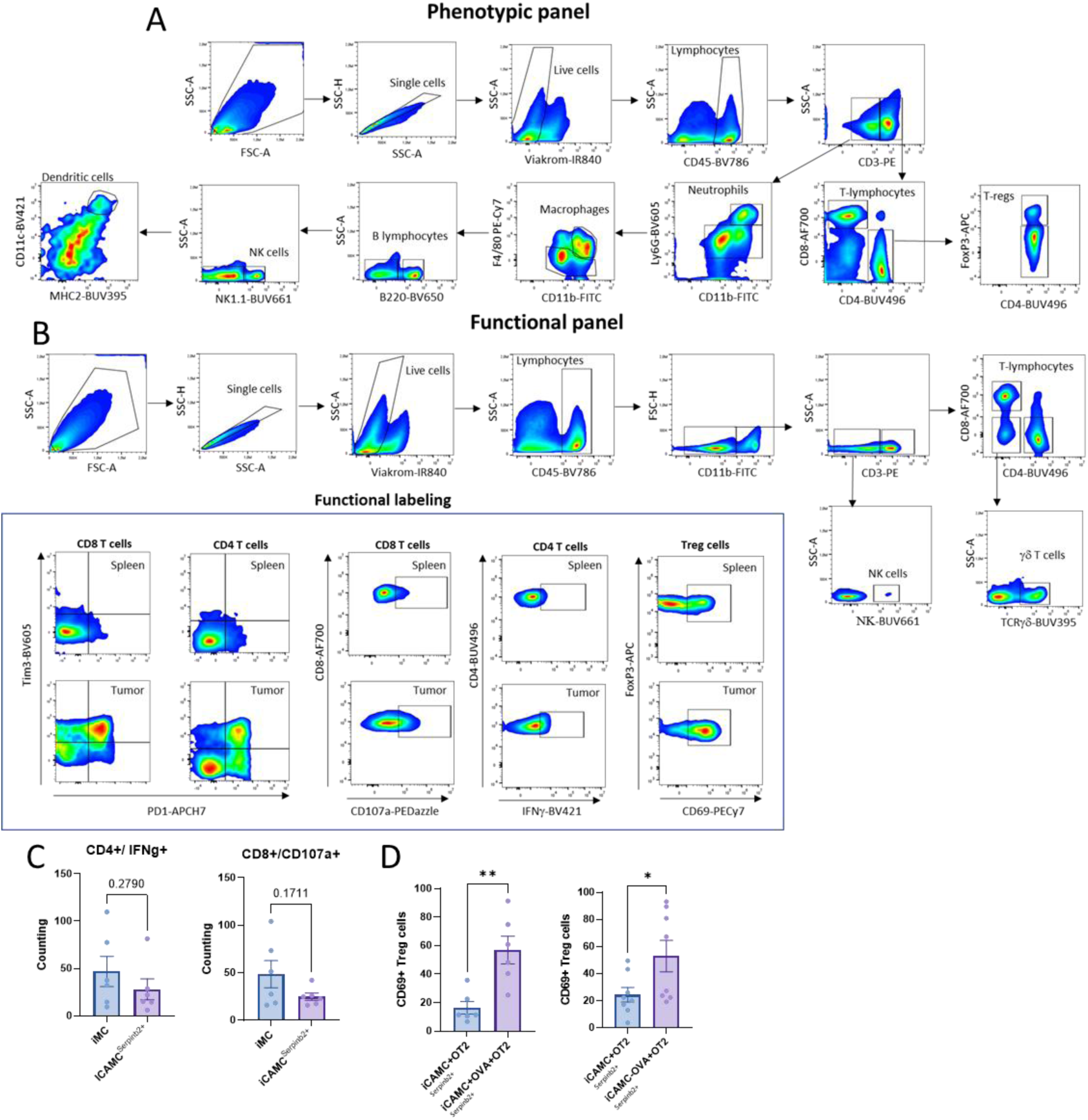
Gating strategy for phenotypic and functional panels. A) Gating strategy of the immune cell panel by flow cytometry to characterize immune cell phenotypes. B) Gating strategy of the immune cell panel by flow cytometry to assess immune cell functions. C) Percentage of CD4^+^ T cells expressing IFNγ and percentage of CD8^+^ T cells expressing CD107a in KPCA tumors mixed with iMCs or iCAMC^Serpinb2+,^ evaluated by flow cytometry. D) Percentage of CD69^+^ cells within the regulatory T cell population evaluated by flow cytometry at 24 h and 72 h.

